# The Translational Landscape of Reactive Astrocytes Reveals the Impact of eIF2B-mediated Dysregulation in VWM Disease

**DOI:** 10.1101/2025.01.14.633074

**Authors:** Shir Mandelboum, Liat Lev-Ari, Andrea Atzmon, Melisa Herrero, Naama Brezner, Daniel Benhalevy, Orna Elroy-Stein

## Abstract

A devastating genetic recessive neurodegenerative disorder, Vanishing White Matter Disease (VWMD), stems from mutations in eIF2B—a master regulator of mRNA translation initiation and mediator of cellular stress response. While astrocytes, the brain’s essential support cells, are known to be central to VWMD pathology, the molecular mechanisms underlying their dysfunction remain poorly understood. Our study reveals that even a mild mutation in eIF2B5 profoundly disrupts astrocyte mRNA translation regulation upon cytokine-mediated activation, affecting nearly one-third of all expressed genes. Through innovative integration of RNA-seq and Ribo-seq analyses using primary cell cultures of astrocytes isolated from eIF2B5^R132H/R132H^ mice, we discovered attempts to compensate for impaired protein production by increasing mRNA levels. However, this compensation proves insufficient to maintain critical cellular functions. Our comprehensive analysis uncovered significant disruptions in cellular energy production and protein synthesis machinery. We also predicted previously unknown defects in cholesterol biosynthesis within mutant astrocytes. Moreover, a meta-analysis of translation initiation scores pinpointed, for the first time, a short list of specific ’effector’ gene candidates that may drive disease progression. This powerful combination of transcriptome and translatome illuminates the complex pathophysiology of VWMD and identifies promising new biomarkers and therapeutic target opportunities.

## Introduction

Regulation of gene expression in response to a physiological cue is a sophisticated dynamic process that evolved to optimize cell function by generating the accurate level of each gene product at the precise timing. While regulating mRNA abundance is often more economical, regulating translation enables a more rapid and dynamic response. Global regulation of protein synthesis is generally mediated by the two most prominent initiation factors, eIF4E and eIF2B. eIF4E, the 5’cap binding protein and driver of canonical 5’cap-dependent translation is typically regulated by mTOR, governing cell growth and proliferation (1). In contrast, much of the published information regarding the role of eIF2B as a translation regulator refers to it being a central node in the cellular response to stress conditions, i.e., the integrated stress response (ISR). At each round of translation initiation, eIF2B recycles the inactive eIF2·GDP back to its active eIF2·GTP form, enabling its re-binding to initiator Met-tRNA^Met^ to re-generate the eIF2·GTP·Met-tRNA^Met^ ternary complex (TC). The TC is then delivered to a 40S small ribosomal subunit towards re-establishing a translation pre-initiation complex. In response to multiple stressors, eIF2α is phosphorylated by one of the four eIF2α kinases (PERK, PKR, GCN2, HRI), leading to lower TC levels and global translation attenuation.

Then, when TC levels are low, the translation of mRNAs encoding rescue proteins is specifically enhanced, promoting cellular ISR. Translation in conditions of low TC is often enabled by a leaky scanning/re-initiation mechanism related to upstream AUG codons (uAUGs)- or upstream open reading frames (uORFs)-burdened 5’-untranslated regions (5’UTRs). Therefore, the regulatory power of eIF2B relies on highly regulated TC levels at each time point, combined with the features of gene-specific 5’UTRs (2–4).

More than 160 partial-loss-of-function mutations in any of the five genes encoding the five subunits of eIF2B result in an eIF2B-leukodystrophy termed Vanishing White Matter Disease (VWMD), a rare autosomal recessive neurodegenerative disorder (OMIM#306896) (5). VWMD is manifested by the gradual depletion of brain white matter (e.g., myelinated axons and surrounding glial cells), resulting in severe neurological symptoms that progressively deteriorate towards death around the early teens.

Although a partial loss of eIF2B function, essential to all cell types, causes dysregulation of general translation, the disease is mainly confined to brain glial cells (6). Astrocytes, the brain ‘homeostasis keepers,’ are known to be central to VWMD (7–9).

Our previous study resolved cell type-specific technical barriers to accomplish a first-ever ribosome foot-printing (Ribo-seq) in primary astrocytes and provided genome-wide information on astrocyte gene expression strategies along cytokine-mediated activation (10). We identified functional gene sets responding by different strategies, e.g., shifting mRNA levels, translation efficiency per se, or both. 30% of expressed genes responded to cytokines by changes in translation efficiency, indicating this is an essential regulatory strategy required for optimal cellular response. In this study, we used our unique capability to assess ribosome density (RD) along primary astrocytes mRNAs to test the effect of a mild loss-of-function of eIF2B. We used the hypomorphic mutation R132H in the catalytic subunit, which leads to a 20% reduction in brain eIF2B enzymatic activity, as implicated in our mouse model for WMVD (11).

Primary astrocytes isolated from WT and eIF2B5^R132H/R132H^ mice were employed to characterize each transcript’s differential mRNA expression and RD, in response to cytokine exposure. Initiation score (IS), reflecting the ratio between translation initiation and elongation rates per transcript, was compiled and used to define a short list of 21 ’effector’ genes responsible for key cellular functions whose translation initiation is the most sensitive to eIF2B mutation. Integrated analysis of shifts in IS and RD revealed insights into the specific expression networks in primary astrocytes that are hyper-susceptible to eIF2B function, offering new insight into WMVD pathophysiology and potential therapeutic strategies.

## Materials and Methods

### Biological resources

Wild type (WT; C57BL strain) and eIF2B5^R132H/R132H^ mutant (Mut) mice were bred and housed in Tel Aviv University animal facility with 14/10 h light/dark cycle in groups of two animals per cage in individually ventilated cages (Lab Products Inc., Seaford, DE, USA) supplemented with autoclaved wood chips.

Animals were fed with autoclaved rodent pellet (1318 M, Hybridpellet, Altromin International) and sterile water ab libitum. All experimental procedures were approved by the Tel Aviv University Animal Care Committee according to national guidelines (permit #04-21-020). Breeding and genotyping were performed as previously described (11). Primary astrocytes were isolated from P0-P2 newborn brains following papain dissociation to single-cell suspension using MACS Neural Tissue Dissociation kit (#130-092-628; Miltenyi Biotec, Bergisch Gladbach, Germany) and GentleMACS dissociator according to the manufacturer’s protocol. Astrocytes were positively selected using anti-ACSA2 MicroBead Kit (#130-097-678; Miltenyi Biotec), seeded at a density of 5 × 10^5^ cells per 6-cm plate in ‘astrocyte medium’ (#1801; ScienCell Research Labs, Carlsbad, CA, USA) until ∼95% confluency (7–10 days, with medium change every 3 days), split by transferring to a 10-cm plate until ∼95% confluency (∼3 days), then to 15-cm plate (∼3 days), then frozen at a density of 10^6^ cells per cryogenic tube and stored in liquid nitrogen. All experiments were performed using freshly thawed primary astrocytes at the above-detailed passage #3, grown in ‘astrocyte medium’. For cytokine-mediated activation, thawed cells were incubated for 72-96 h (depending on confluency) followed by treatment with 20ng/ml TNFα (Peprotech #315-01A) and 0.25 ng/ml IL-1β (Peprotech #211–11B) for 24 h or 48 h.

### Polysome profiling and detection

WT and Mut Primary astrocytes were seeded at a density of 1.3 × 10^6^ cells per 15-cm plate in ‘astrocyte medium’ and incubated for 120 h until they reached a density of ∼2.5 × 10^6^ cells per plate. Four 15-cm plates of primary astrocytes were used per single sucrose gradient tube. Cycloheximide (CHX) was added to the growing medium at a final concentration of 100 μg/ml for the last 5 min of incubation, followed by medium removal and gentle wash with ice-cold PBS, on ice. A volume of 230 µl per plate of lysis buffer, composed of 25 mM K-HEPES pH 7.2, 10 mM MgCl_2_, 200 mM KCl, and freshly added 2% n-Dodecyl β-D-maltoside (DDM), 1 mM DTT, 1× complete EDTA-free protease inhibitor (Roche), 100 μg/ml cycloheximide, and 1 U/μl RiboLock RNase inhibitor (Thermo Scientific), was dripped directly onto the cells monolayer on ice, followed by gentle spreading to cover the entire plate surface, for 5 min incubation on ice. The scraped, collected and combined lysed cells were incubated on ice for an additional 10 min, followed by centrifugation at 2,000 × g for 5 min at 4°C for nuclei removal. The supernatant was centrifuged at 20,000 × g for 10 min at 4°C for membranes and organelles removal. The entire quantity of recovered supernatants yielded from 4x15-cm plates was then loaded onto 9 ml of 15.7–50% gradients in 13.2 ml polypropylene tubes (Beckman coulter 331372) and centrifugated at 37,000 rpm for 90 min at 4°C using SW41 Beckman rotor. Gradients were fractionated using a Teledyne ISCO UA-6 UV/VIS gradient elution and detection system. Polysome profiles were obtained through continuous absorbance measurement at 254 nm. 18–20 fractions (0.6 ml each) representing the entire gradient were collected into tubes containing SDS at a final concentration of 1%.

### Ribosome foot printing (Ribo-seq), RNA-seq, and analysis

Three biological repetitions of WT and Mut primary astrocytes that had been treated or not with cytokines for 24 and 48 h were used for ribosome foot printing as detailed previously (12). Cells were lysed as described above for polysome profiling, with two adjustments: (i) Prior to harvesting, the cells were incubated with 2 μg/ml of Homo-Harringtonine (Tocris #1416) for 2 min at 37°C, to enrich for translation initiation complexes; and (ii) the lysis buffer lacked the RNase inhibitor component and included 25 U/ml Turbo DNase-I (Thermo Scientific, AM2238). A portion of each sample was treated with RNase-I (Thermo Scientific, AM2295) to generate ribosome footprints, while an undigested portion was used to isolate total RNA which was then poly(A) selected using NEBNext® Poly(A) mRNA Magnetic Isolation Module (New England Biolabs, #E7490). Following RNase treatment, the footprint samples were layered on 1M sucrose cushion, following ultracentrifugation at 78,000 rpm for 2 h at 4°C using a Beckman Coulter TLA120.2 rotor. RNA extracted from sucrose cushion pellets were subjected to size selection, by separation of the footprint samples on a 15% polyacrylamide TBE–urea gel followed by purification of 26–34 nt fragments from the gel. rRNA depletion was achieved using Ribo-Zero Gold, then libraries were generated with NEBNext Multiplex Small RNA Library Prep Set for Illumina (New England Biolabs, #E7300L). Next, libraries were size selected by gel electrophoresis on 8% TBE-PolyAcrylamide gel, with ∼155 bp sized product purified from the gel. For Poly(A)-selected RNA, libraries were generated using TruSeq Stranded Total RNA Library Prep Gold (Illumina #20020598). For both Ribo-seq and RNA-seq, sequencing was performed using Illumina NextSeq 550 platform using the High Output Kit v2.5 (Illumina #20024906).

Data Analysis: Sequenced reads were aligned to mm10 reference genome with TopHat2 (13), followed by generation of gene count data using FeatureCounts (14). Only genes with at least 20 counts in all replicate samples of at least one condition were included in the analysis. Quality control of Ribo-seq data was carried out using Trips-viz (15) for analyzing RPF length distribution, triplet periodicity, and the distribution of RPF on transcript regions. Conditional quantile normalization (CQN) was performed to correct for sample-specific gene length effect (16). Cqn normalized expression values were clustered using Morpheus (https://software.broadinstitute.org/morpheus). The maximum and minimum values were used to convert all relative values to colors. Kmeans clustering was performed on all genes (rows) using 1 minus Pearson correlation with *K* = 6 clusters. Changes in Ribosome Density (RD) of the coupled Ribo-seq and RNA-seq data were calculated using DTEG analysis (17). GSEA was performed on the processed data with the pre-ranked GSEA method (GSEAPreranked function; GSEA v4.2.3), testing enrichment for KEGG and GO gene sets. Next, log_2_ fold change of specific subsets of genes (selected among the significant GSEA results) was compared with the expression of all the rest of the genes in the dataset (which served as background). *P*-values were calculated using Wilcoxon’s test. All log2 fold-change values and corresponding p-values are provided in Supplementary Table S1. Relative expression maps of RNA-seq and Ribo-seq were generated with ‘Proteomaps’ (18). Polygon area was quantified using SketchAndCalc™ Area Calculator (http://sketchandcalc.com Vers 4.1.3).

### Composition of ‘ribosomal proteins’, ’OXPHOS’ and ‘cholesterol biosynthesis’ gene subsets

The ‘ribosomal proteins’ (RP) subset includes the 79 RP genes (19) (of which only 77 are expressed in our data). ‘OXPHOS’ is GO:0006119, n=75. The ‘cholesterol biosynthesis’ subset (n==30) includes all enzymes of the mevalonate, Bloch, and Kandutsch-Russell pathways and key regulators of cholesterol biosynthesis, as follows: HMGCR, HMGCS2, IDI1, FDFT1, SQLE, LSS, ACAT1, ACAT2, IDI2, GGPPS, CYP51, NSDHL, SC5D, DHCR7, HSD17B7, LBR, MSMO1, INSIG1, INSIG2, **MVK, PMVK, FDPS, MVD, EBP, DHCR24**, **SCAP, SREBF1, SREBF2** (Figure 4, ‘Translationally-regulated’ are in bold). ACAT3 and HMGCS1 (claimed as ‘predicted’) were highly expressed in our data and thus were included too. To classify the subsets into ’non-translationally-regulated’ and ’translationally-regulated’, we used the cutoffs of WT RD(24hr)>0.3, WT(24hr)>0.1, or WT(48hr)>0.3 for RP, OXPHOS and cholesterol biosynthesis, respectively.

### Relative Ribosome Occupancy (RRO) and Initiation Score (IS) calculations

Single nucleotide resolution ribosome occupancy was generated using RiboTricer 1.4.0 (20), using gencode_vM25_annotation_mm10.gtf ORF data. RiboTricer default settings were employed, using AUG-start codons for ORFs detection. Before downstream analyses, data was filtered to include: (i) Only annotated ORFs. (ii) A single transcript per gene was selected based on the transcript with the highest read density in the WT (0 hr). Replicate B was selected as it had the highest gene coverage. These transcripts were then matched across all other samples. Relative Ribosome Occupancy per nucleotide was calculated as [10Xreads per position], divided by [sum of reads per ORF excluding the first 30 nucleotides]x[ORF_Length-30]. For each replicate, the Initiation Score (IS) per gene was calculated as the ratio between start codon reads density (sum of nucleotides in positions 1-3) divided by 3, and elongation reads density (defined as the sum of reads excluding the first and last 30 reads of the ORF, divided by the length of the ORF minus 60). IS values per gene were averaged across replicates after confirming strong correlations between replicates. Genes lacking IS values in all four conditions (WT/Mut; 0 hr/24 hr) were excluded, resulting in a final dataset of 9,574 genes. For the selection of IS-regulated genes (see Figure 5), the dataset was further filtered using the following criteria: (i) Elongation Reads density > 0.1 (excluding extremely low detected genes); (ii) The sum of IS scores across all four conditions > 1 (ensuring sufficient signal for comparative analysis). This filtering yielded a final dataset of 2,332 genes for IS analysis. To judge condition-dependent alterations in IS values, the fold-changes (FC) values were classified into response categories: upregulation, mild up, no response, mild down, and downregulation. Uncalculatable FC values due to IS=0 in the numerator or denominator were attributed to a category based on absolute IS values.

### Measurement of total cell cholesterol

Three biological replicates of WT and Mut astrocytes were seeded at 7.5 × 10^4^ cells per well of a 6-well plate and incubated for 120 hours, with or without 20 ng/ml TNFα and 0.25 ng/ml IL-1β during the final 48 hours. Following washing with PBS, lipids were extracted from cell monolayers by adding 300 μL of a ’Lipid extraction mix’ (chloroform:isopropanol:IGEPAL-CA-630 at a ratio of 7:11:0.1) per well, followed by scraping and content collection. Cholesterol level was quantified enzymatically using a Cholesterol/Cholesteryl Ester Quantitation Kit (Biovision, K603-100) as detailed in the manufacturer’s protocol. The cholesterol signal was measured using a fluorescent plate reader (SYNERGY H1, Biotek) with excitation/emission set at 535/590 nm, using extended gain. Cholesterol values were normalized to protein content, as determined by using Pierce™ BCA Protein Assay Kit (Thermo-scientific #23225) per manufacturer’s instructions.

### Oxidative respiration assay

WT and Mut Primary astrocytes were seeded in ‘astrocyte medium’ at a density of 5 x10^3^ cells per well of XF-96-well microplate pre-coated over-night with 10 ug/mL of Poly-D-Lysine (PDL) (#P0899; Sigma-Aldrich). The cells were incubated for 72 h at 37°C with or without 20 ng/ml TNFα and 0.25 ng/ml IL-1β during the last 48 hr of incubation. Oxygen Consumption Rate (OCR) for basal and maximal mitochondrial respiration was determined as previously described (21).

### Statistical analyses

Transcriptome and translatome libraries were prepared and sequenced following ENCODE Experimental Guidelines for ENCODE3 RNA-seq. For each experiment performed with astrocytes, ≥3 biological repeats were used, each containing a different batch of astrocytes isolated from independent litters. Unless otherwise stated, statistical analysis was performed with Prism 9.0.1 software (GraphPad). For RNA-seq/Ribo-seq data, Wilcoxon test was used for comparing ‘GO’ and ‘KEGG’ subsets, or manually defined subsets of genes, to all the rest of the genes in the dataset, using R software (v3.5.1) (R-project.org) and RStudio (rstudio.com). P-values were determined by two-way ANOVA and Sidak’s multiple comparison test for the oxidative respiration assay. For RRO analysis, GraphPad software (Prism version 10.4) was used for plotting either Scatter plot with Bar with SEM or Box and whiskers plot (Tukey); and statistical analysis for IS distribution in cluster #2 vs. cluster #4 Kolmogorov-Smirnov test, and any comparisons of the four conditions (WT/Mut; 0hr/24hr) 2-way-ANOVA.

## Results

### A hypomorphic mutation in eIF2B5 disrupts gene expression programs along cytokine-mediated activation of astrocytes

To closely examine how changes in gene expression are executed in eIF2B-mutant astrocytes, we used cultures of primary astrocytes isolated from WT and eIF2B5^R132H/R132H^ (Mut) mice (11). Biological replicates were treated or not with TNFα and IL1β for 24 or 48 hr, followed by RNA-seq and Ribo-seq analyses. Given the pathogenic nature of the R132H mutation in the catalytic subunit, we hypothesized that slightly lower ternary complex (TC) concentration due to the 20% less active eIF2B would reduce cellular capacity to regulate translation initiation. To enhance detection, we combined translation initiation and elongation inhibitors into the Ribo-seq procedure to stabilize initiating and elongating ribosomes onto their mRNA templates. We have used Harringtonine conditions that allow quantifying differential Mut versus WT translation initiation events. Ribo-seq quality was confirmed (**Supp. Figure S1**), with RPF length distribution at 25-30 nt long and predominant alignment to mRNAs coding sequence (CDS) at triplet periodicity.

Principal component analysis (PCA) revealed WT and Mut separation at PC1, PC2, and PC3 (**Figure 1A**: RNA-seq, left panel; Ribo-seq, right panel) and treatment status separation at PC1, showing a clear distinction between untreated and cytokine-treated samples regardless of their genotype (PC1 RNA-seq: 69%; PC1 Ribo-seq: 64%). Reduced separation between WT and Mut samples based on RNA-seq data, relative to their greater separation based on Ribo-seq data, suggests the two genotypes apply a relatively similar transcriptional response to cytokines and indicates that the eIF2B mutation predominantly impacts translation. RNA-seq and Ribo-seq data of untreated astrocytes and following 24 and 48 hr of cytokines exposure was used to cluster 11,449 expressed genes into six groups representing different expression strategies, emphasizing WT vs Mut similarities and dissimilarities (**Figure 1B**). Most genes exhibited changes in mRNA abundance followed by congruent translation without a change in their RD. In WT astrocytes, 68% of the genes respond this way. Specifically, 38% were downregulated (cluster #1), 12% were gradually and consistently upregulated, reaching maximal values 48 hr after cytokines treatment (cluster #2), 8% were transiently upregulated at 24 h followed by a decrease at 48h (cluster #3), and 10% were upregulated with limited congruent translation (cluster #5) (**Figure 1B**, pink and grey boxes). The remaining genes exhibited expression response strategies based on mRNA translation regulation. In WT, 32% of the genes were regulated this way, where 18% were translationally upregulated (cluster #4) and 14% were translationally downregulated (cluster #6) (**Figure 1B**, two light blue boxes). Our downstream analysis focused on genes in clusters #2 and #4. Both clusters include genes upregulated in response to cytokines in non-facultative strategies, with cluster#2 genes exclusive mRNA upregulation and cluster#4 genes exclusive RPF upregulation. As shown, while mild WT vs Mut differential responses are observed in all clusters, clusters #4 and #6, which include genes responding at the translational level, exhibit higher differences in WT vs Mut astrocytes relative to other clusters.

**Figure 1.**
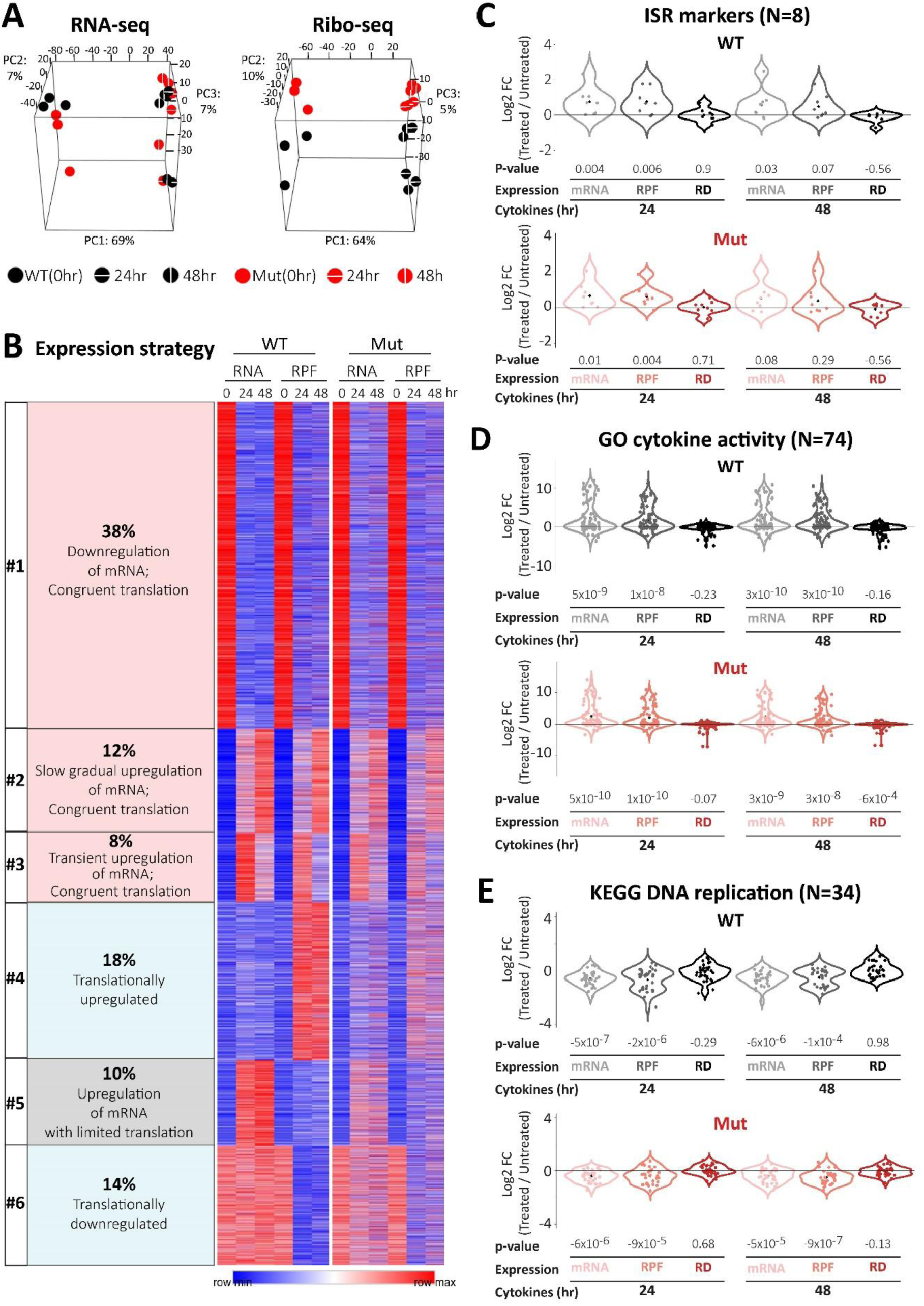
A point mutation in eIF2B adversely affects mRNA translation regulation programs during cytokine-mediated activation of astrocytes. WT and eIF2B-mutant (Mut) astrocytes were treated or not with TNFα (20 ng/ml) and IL1β (0.25 ng/ml) for 24 and 48 hr followed by RNA-seq and Ribo-seq analyses. (**A**) Principal component analysis (PCA) applied to the RNA-seq (left) and Ribo-seq (right) data. Three biological replicates are presented for each sample. WT, black; Mut, red; 24 hr, horizontal white line; 48 hr, vertical white line. Shown is a scatter plot of all replicates for the primary three principal components (PC1, PC2, and PC3). (**B)** Heatmaps of cqn normalized expression values. K-means clustering (K=6) was applied to all genes (N=11,449; rows). (**C-E)** Log2 fold-change (FC) values of total mRNA level, RPF, and RD of individual gene products obtained in response to 24 and 48 hr cytokine treatment relative to untreated WT and eIF2B-mutated astrocytes. Three gene subsets are presented: **(C)** ISR markers upregulated in astrocytes upon ER stress, detailed in Table 1 of Sims et al., 2022 (N=8). **(D)** ‘GO cytokine activity’ (N=74). check(**E**) ‘KEGG DNA Replication’ (N=34). P-values were calculated using Wilcoxon’s test, comparing the selected gene sets to the rest of the genes in the dataset.

Since activation of the ISR is wired through the PERK arm of the unfolded protein response (UPR), which accompanies activation of astrocytes by cytokines (22), we wanted to confirm our experimental conditions recapitulates upregulation of ISR markers related to ER stress. Cytokine exposure led to higher expression of most ISR markers in both genotypes, without a significant difference between WT and Mut astrocytes (**Figure 1C**). Some genes were transiently upregulated at the mRNA level followed by congruent translation (**Figure 1B** cluster #3, e.g., ddit3, Ppp1r15a, Trib3), others were upregulated at the mRNA level with limited translation (**Figure 1B** cluster #5, e.g. Atf6b, Ero1lb, Gadd45a), and ATF4 was upregulated at the translation level (**Figure 1B**, cluster #4). The similar induction of ISR markers in WT and Mut astrocytes is inconsistent with a direct link between the hypomorphic eIF2B mutation and ISR, implying the possibility that eIF2B5^R132H/R132H^-driven dysregulation is ISR-independent.

We previously showed that during astrocytes’ response to cytokines, functionally related genes share common expression strategies (10). To assess the impact of eIF2B mutation on a translational response, we first wanted to rule out an effect on the modulation of mRNA levels. As shown, Mut astrocytes recapitulated mRNA levels upregulation, with no significant change in RD, for cytokine-related genes (**Figure 1D**; N=74; P ≤ 1.00E-08, Supp Table 1) and apoptosis-related genes (**Supp Figure S2**, N=67; P ≤ 1.00E-05). Similarly, Mut astrocytes recapitulated the downregulation of mRNAs encoding DNA replication genes (**Figure 1E**; N=34; P ≤ 1.00E-04). These results exclude a direct link between mutated eIF2B and mRNA abundance dysregulation. Therefore, we transitioned to test the straightforward hypothesis that eIF2B mutation interferes with astrocytes’ capacity to modulate translation in response to acute demands.

### eIF2B mutation negatively impacts the regulation of specific gene sets

Given that ribosomal proteins (RPs) mRNAs are known to be tightly translationally regulated (23), we tested whether the mutated eIF2B compromises this regulation. Our data enabled the classification of the 77 RP encoding mRNAs (19) into two categories: 30% (n=23) were not translationally regulated, while 70% (n=54) were translationally regulated. Inspection of the first group revealed similar expression across genotypes, demonstrating increased mRNA abundance and unchanged ribosome density (RD) 24 hr after cytokines treatment **(Figure 2A**, upper panel). However, a differential WT vs Mut response was observed for the second sub-group of transcripts. In WT cells, 40 of the 54 translationally regulated RP genes responded to cytokines by an increase in RD driven by the combined reduction of mRNA levels and an increase in RPF (all 40 genes reside in cluster #4, Figure 1B). In contrast, the RD of these genes in Mut astrocytes could not be upregulated, and the response often included mRNA upregulation, possibly as a compensatory mechanism. Noteworthy, in many cases, mRNA upregulation in Mut cells was coupled to a significant decrease in RD values, indicating considerable difficulty in increasing translation. In agreement with a phenotype driven by eIF2B mutation, Mut astrocytes fail to increase ribosomes occupancy near RP transcripts start codons, as observed in WT astrocytes (Representative data for RPL19 and RPL14 is shown in **Figure 2C** and **Supp Figure S3**). To look into the full scale of differential expression responses in WT and Mut astrocytes (**Figure 2B**, upper panel), we compiled relative expression maps by integrating our data with pre-classified functionally related pathways (‘proteomaps’) (18). The generated relative expression maps of WT and Mut astrocytes before and after 24 hr of cytokine treatment are shown in **Supp Figure S4**. Zooming into the ‘Ribosome’ group, which refers to genes encoding the RPs (N=77; red line-surrounded polygon in **Supp Figure S4**), WT cells demonstrate limited translation of the large reservoir of RP-encoding transcripts. While RP transcripts comprise 3.4% of the mRNA population, their RPFs comprise only 0.4% of total RPFs. After Cytokines induction for 24 hr, they increase to only 0.7% of total RPFs, indicating a remarkable reservoir of RP transcripts, which probably enables rapid RP upregulation upon a signal (10,24), and is mildly utilized upon exposure to cytokines. In Mut cells, however, the main response to cytokines was in the form of mRNA levels upregulation, comprising 2.6% of total mRNAs and reaching 3.3% (**Suppl Figure S4C**). This observation demonstrates a seemingly limited capacity of Mut astrocytes to upregulate translation upon demand, coupled with compensatory adaptation mechanisms. The observation that Mut astrocytes maintain a lower RP mRNA reservoir at steady state (76% that of WT) and its build-up to almost WT level upon cytokine exposure hints at the reduced protein expression capacity of Mut already before cytokines induction.

**Figure 2.**
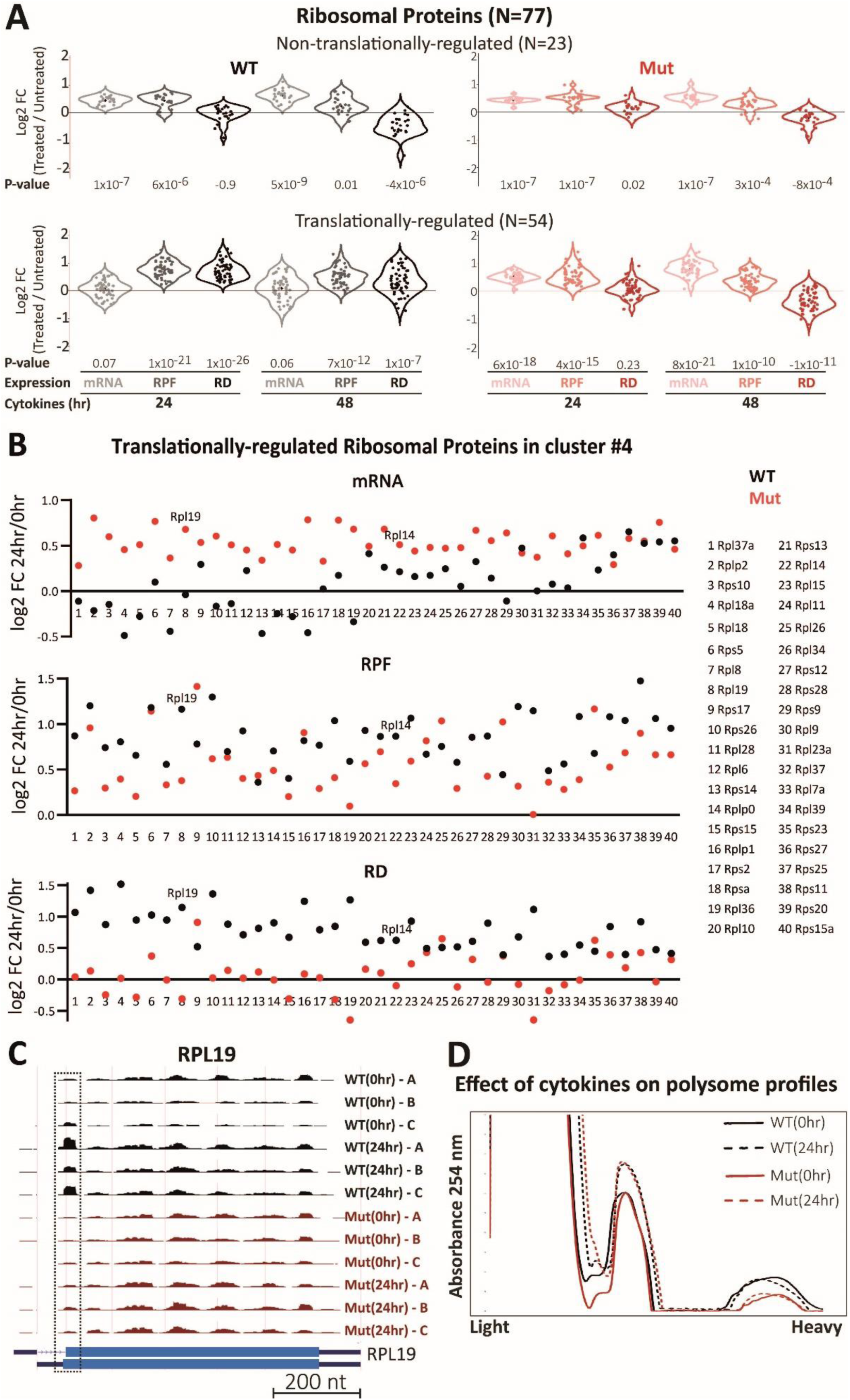
Effect of eIF2B mutation on expression response of genes encoding ribosomal proteins and its functional consequence. **(A)** Transcripts encoding ribosomal proteins (N=77) classified based on their log2FC RD value in WT at 24 hr were parted into two groups: (i) non-translationally regulated (RD<0.3; N=23, top) and (ii) translationally-regulated (RD>0.3; N=54; bottom). Log2 FC values obtained for WT and Mut are shown as detailed in Figure 1(C-E). **(B)** Log2 FC values of mRNA, RPF, and RD of each translationally-regulated RP gene in cluster #4 (Figure 1B), upon 24 hr of cytokine treatment. **(C)** RPF map of RPL19, highlighting the increase in RPF reads at the AUG start codon (dashed rectangle), in WT but not in Mut, in response to 24 hr of cytokine treatment. (**D**) WT and Mut astrocytes were treated or not with cytokines for 24 hr followed by polysome profiling analyses. WT, black; Mut, red; untreated (0 hr), straight line; 24 hr, dashed line.

Polysome profile evaluation of WT and Mut astrocytes under both conditions revealed that both genotypes could equally increase their monosome quantity upon 24 hr of cytokine exposure, suggesting a satisfactory compensatory response by Mut astrocytes (**Figure 2D**). However, a lower number of polyribosomes was evident in Mut before and after cytokine treatment, regardless of the sufficient amount of monosomes. This phenomenon supports the notion of inefficient translation initiation by the generated monosomes in Mut. A similar translation dysregulation was also observed in the Rrn3 gene encoding RNA polymerase 1, which is required for ribogenesis (**Supp Figure S5**, note the increase in RPF reads at the AUG start codon, in WT but not in Mut, in response to cytokines).

### eIF2B-mutant astrocytes fail to execute the translation regulation program of the oxidative phosphorylation gene subset

Our previous studies revealed that energy production by oxidative phosphorylation (OXPHOS) is defective in the brains of eIF2B5^R132H/R132H^ mice and primary fibroblasts and astrocytes cultures generated from this Mut mouse strain (21,25,26). To probe whether translation dysregulation might drive this phenotype, we inspected how genes implicated in OXPHOS (GO:0006119, n=75) are differentially regulated in WT and Mut astrocytes. In WT astrocytes, we identified 48 OXPHOS genes that were translationally regulated in response to cytokine induction and 27 that did not demonstrate a translational shift. All 27 genes that were not translationally regulated exhibited a similar expression response in Mut, with similar mRNA levels and RD values (**Figure 3A**, upper panel). However, genes that showed a translational response in WT, failed to respond in Mut astrocytes. Specifically, in WT, these genes exhibited an increase in RD, resulting from a combined decrease in mRNA abundance and an increase in RPF, while Mut astrocytes seemingly compensated by a lack of decline in mRNA abundance to facilitate a sufficient level of reads without upregulating translation rate. The data from Mut astrocytes indicates a further worsening of translation ability at 48 hr post cytokines induction, showing a more significant reduction in RD coupled with a further increase in mRNA abundance (**Figure 3A**, lower panel; p=-0.01). Specific examples include Ndufa11, Uqcrq, and Atp5d, which represent mitochondrial membrane-bound electron transfer complex (ETC) I, III, and V subunits, respectively. The translationally regulated OXPHOS genes are enriched in cluster #4, but due to their variable sensitivity to eIF2B mutation, some reside in cluster #1 (**Figure 1B**). This variability, even within cluster #4, is emphasized in **Figure 3B**, which shows the fluctuating RD values in WT because of the fluctuated decrease in mRNA abundance. In contrast to the general trend, Cox6c, a subunit of the electron transfer complex IV, remained similarly regulated in Mut astrocytes, with comparable mRNA abundance, RPF, and RD (**Figure 3B**). Inspection of the distribution of RPF along the Cox6c transcript revealed an uORFs-mediated translation regulation, providing a classic example of how RD values may be misleading in cases where translation can be differentially partitioned between uORFs and the main ORF (**Figure 3C**). Specifically, in WT astrocytes RPF along the Cox6c transcript demonstrated efficient initiation at the uORF and attenuated initiation at the main ORF in response to cytokines. However, leaky scanning through the uORF increased ribosome occupancy along the main ORF in Mut astrocytes (**Figure 3C**). Interestingly, ribosomes seemed to accumulate downstream of the transmembrane domain coding region, possibly to allow precise protein folding/membrane integration/co-assembly with other subunits. This observation and uORF regulation is further discussed later.

**Figure 3.**
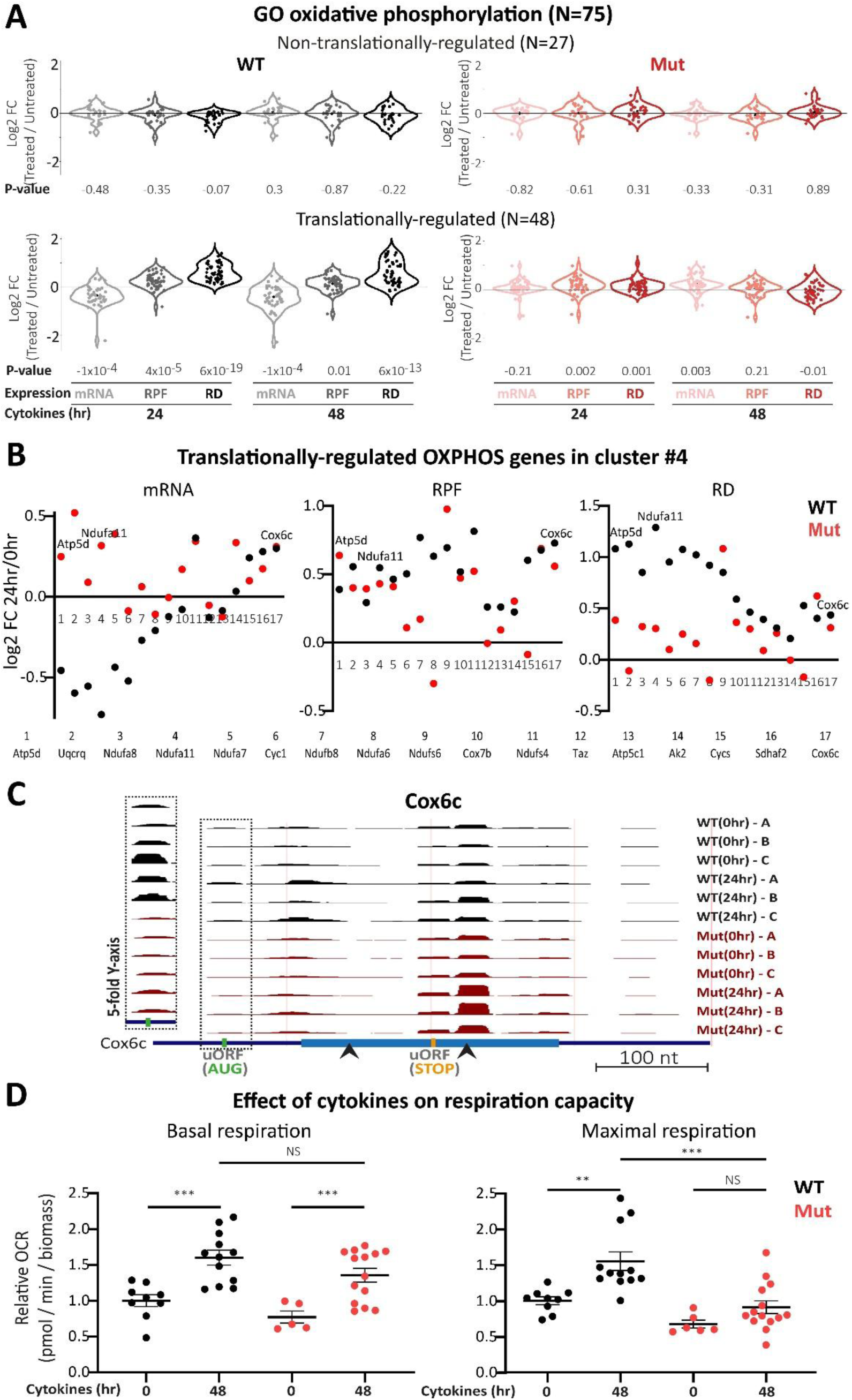
Effect of eIF2B mutation on expression response of genes related to OXPHOS and its functional consequence. **(A)** Members of the ‘GO oxidative phosphorylation’ subset (N=75) classified based on their Log2FC RD value in WT at 24 hr were parted into two groups: (i) non-translationally regulated (RD<0.1; N=27; top), and (ii) translationally-regulated (RD>0.1; N=48; bottom). Log2 FC values obtained for WT and Mut are shown as detailed in Figure 1(C-E). (**B**) Log2 FC values of mRNA, RPF, and RD of each translationally-regulated OXPHOS gene that reside in cluster #4 (Figure 1B), in response to 24 hr of cytokine treatment. **(C)** RPF map of Cox6c, highlighting the increase in RPF reads at the uORF (dashed rectangle) in WT but not in Mut, in response to 24 hr of cytokine treatment. uORF start codon, green; uORF stop codon, orange. Black arrows mark the sequence encoding the trans-membrane domain. Note the increased ribosome accumulation at the C-terminus of the transmembrane domain in Mut compared to WT. **(D)** WT (black) and Mut (red) astrocytes were treated or not with cytokines for 48 hr and subjected to oxidative phosphorylation analysis. Values of oxygen consumption rate (OCR) normalized to the untreated WT values are presented for basal respiration (left) and maximal respiration (right). The average values ± SEM of 5–15 replicates of two independent experiments are shown. P-values were determined by two-way ANOVA using Sidak correction for multiple comparisons. *P ≤ 0.03; **P ≤ 0.002; ***P ≤ 0.0004; NS, not significant.

We compared oxygen consumption rates in Mut and WT astrocytes to test whether the compromised expression response due to the hypomorphic mutation in eIF2B affected OXPHOS capacity. A ∼1.5-fold increase in both basal and maximal respiration following cytokine exposure was observed in WT astrocytes. Mut astrocytes exhibited a similar increase in basal respiration. However, maximal respiration did not increase in Mut, remaining at <60% of the WT level (**Figure 3D**). This result demonstrates how eIF2B-mediated regulation might contribute to an adaptive cell response to cytokines. Overall, our analyses highlight a reduced capacity of Mut astrocytes to regulate OXPHOS gene translation in response to cytokines, which is partially compensated by alternative expression mechanisms and presents a compromised energy metabolism.

### Cholesterol biosynthesis is sensitive to eIF2B mutation - a new insight into VWMD pathology

Next, we set out to test the ability of our data to provide meaningful molecular hints that may explain additional phenotypes associated with VWMD pathology. A key clinical feature is the loss of brain white matter due to impaired maintenance of myelin (6,27). The vital connection between cholesterol biosynthesis in astrocytes and white matter regeneration in the brain (28) prompted us to inspect the involved genes in-depth. For this purpose, we defined a ‘cholesterol biosynthesis’ subset that includes genes encoding enzymes of the mevalonate pathway, the Bloch/Kandutsch-Russell pathways, and seven key regulators (n=30, of which 28 were detected in our data). We then dissected the ‘cholesterol biosynthesis’ subset into 19 non-translationally regulated, and nine translationally regulated genes. The expression response of non-translationally regulated genes was similar across genotypes, with mRNA upregulation at 24 hr post cytokines treatment (WT, p=0.0003; Mut, p= 0.005) followed by a congruent increase in RPF (**Figure 4A**, upper panel; WT, p=0.01; Mut, p=0.02). These included FDFT1, a rate-limiting enzyme in the mevalonate pathway branchpoint towards cholesterol biosynthesis, which is known to be transcriptionally upregulated by SREBF1 and SREBF2, major transcription factors of sterol biosynthesis (29). WT and Mut astrocytes showed a similar increase in FDFT1 mRNA and RPF levels, with hardly changed RD values (**Figure 4A**, upper panel). In contrast, the expression response of the translationally regulated genes was not recapitulated in Mut astrocytes. WT cells presented a significant increase in RD at 24 and 48 hr (p=0.0004 and p=0.0001, respectively), with gene-specific strategies based on changes in RPF and/or mRNA values. However, in Mut astrocytes, the RD values of these genes did not increase at either time point (**Figure 4A**, bottom panel). The differential response of the nine translationally regulated genes at 24 hr post cytokines treatment is demonstrated in **Figure 4B**. As shown, Mut astrocytes presented mixed deficiencies driven by the inability to raise RPF values (e.g., Mvd, Fdps, Ebp) or to lower mRNA levels (e.g., Mvk, Srebf1, Scap, Pmvk). These observations point to a differential expression response across genotypes.

To test whether the eIF2B mutated genotype presents altered functionality, we measured total cholesterol levels in WT and Mut astrocytes before and after cytokine treatment. Our experiment revealed that WT astrocytes exhibit a 1.25-fold increase in total cholesterol in response to cytokine-mediated activation. However, Mut astrocytes failed to increase cholesterol levels (**Figure 4C**). This observation suggests that WT levels of eIF2B enzymatic activity in astrocytes enable the translational fine-tuning required for appropriate cholesterol biosynthesis in response to demand.

### Genes initiation score (IS) as a predictor for eIF2B-dependent regulatory capacity

An important goal of the current study was to pinpoint dysregulation driven by eIF2B mutation, which eventually causes VWMD. The observed broad dysregulation, comprising transcription, mRNA turnover, and translation programs, cannot be directly attributed only to translation initiation dysregulation by mutant eIF2B. Therefore, we hypothesized that a small group of genes, hyper-sensitive to aberrant TC levels driven by eIF2B mutation, encode critical cellular functions that indirectly drive downstream widespread dysregulation. Even though the RD value is widely regarded as a measure of active translation, it does not reliably report initiation efficiency because the spread of RPF reads along the entire coding sequence is dominantly affected by translation elongation dynamics. RD values would also not detect uORFs-mediated regulation. Hence, to focus on initiation events, we first used RiboTricer, to compute the relative ribosome occupancy (RRO) per each nucleotide along the CDS (**Supp Figure S7**) and then calculated initiation scores (IS), representing a standardized initiation/elongation ratio per gene. The IS values were calculated per genotype under each condition, followed by analyzing how they shift between samples by determining their fold change (FC) per gene. IS FC values in response to cytokines treatment (24hr/0hr) revealed that in WT astrocytes, the IS value of 538 genes is upregulated (FC>1.5), and the IS value of 388 genes is downregulated (FC<0.5). However, in Mut astrocytes, the IS of fewer genes is upregulated (N=271), and more genes present IS downregulation (N=448) **(Figure 5A, 5B**). To further compare the IS response to cytokines in WT and Mut astrocytes, we crossed the lists of responding genes and found that only 79 genes were upregulated in both genotypes. Similarly, in response to cytokines, WT and Mut astrocytes shared only 69 downregulated genes. Thus, most genes presenting an IS response to cytokines differ between the genotypes **(Figure 5A, 5B**). This observation suggests that the hypomorphic mutation in eIF2B causes translation initiation dysregulation of hundreds of genes upon cytokine exposure. To further test the relevance of IS value as a predictor for translation regulation capacity, we sought to analyze how IS values of untreated WT astrocytes distribute between cytokine-responsive genes grouped by distinct regulatory strategies. To this end, we focused on clusters #2 and #4 (**Figure 1B**), comprising genes that respond to cytokine treatment in exclusive yet distinct ways. While cluster #2 genes respond exclusively by shifts in mRNA abundance, genes in cluster #4 respond exclusively by shifting ribosome occupancy. In support of the value of IS as a predictor for translation regulation capacity, we found a wider distribution and a 1.5-fold higher IS average value in cluster #4 compared to cluster #2 (p=0.01, **Figure 5C**). To test a link between the IS feature and a Mut phenotype, we observed how IS values differ between each of the WT(0hr), WT(24hr), Mut(0hr), and Mut(24hr) datasets. We first focused on genes with relatively low IS values in WT(0hr) (e.g., 0<IS≤10). While in WT astrocytes, a significant number of these genes exhibited a rise in their low IS value upon exposure to cytokines (IS_WT24hr_ vs IS_WT0hr_, p<0.0001, **Figure 5D**, left), in Mut they failed to increase above the low values, which decreased even further (IS_Mut24hr_ vs IS_Mut0hr_ and IS_WT24hr_ vs IS_Mut24hr_ p<0.0001, **Figure 5D**, left). This observation indicates the limited capacity of Mut astrocytes to modulate translation initiation. Looking at genes with a relatively high IS value in WT(0hr) astrocytes (e.g., IS>10), we noticed that already in the absence of cytokines, these genes in Mut astrocytes are characterized by significantly lower IS values, aligning with the hypothesis that they suffer from a reduced capacity to regulate translation of these genes (IS_Mut24hr_ vs IS_Mut0hr_, **Figure 5D**, right). Accordingly, while the high IS values demonstrated a decline in WT astrocytes in response to cytokines, they showed no sensitivity to the treatment in Mut astrocytes as they were already downregulated before the treatment (IS_WT24hr_ vs IS_WT0hr_ p<0.0001; compared to IS_Mut24hr_ vs IS_Mut0hr_, **Figure 5D**, right). Notably, although the IS values of these genes in Mut astrocytes are much lower than in WT, they did not respond by an increase in IS upon cytokines treatment (p<0.0001, **Figure 5D**, right).

We defined the genes whose IS are uniquely upregulated by cytokine-mediated activation in WT (FC>1.5) but not in Mut astrocytes (**Figure 5B, N**=459). Of those, only 14.6% were able to be slightly upregulated in Mut (1.2<FC≤1.5), while for the rest of them, IS values were either not changed (32%), mildly downregulated (29.6%), or strongly (FC<0.5) downregulated (23.8%, 109 genes, **Figure 6A**, left). These 109 genes, which responded to cytokines in opposite trends depending on eIF2B5(R132H) mutation (**Figure 6A**, dark blue category), included 21 genes that we found to respond to cytokine activation exclusively at the translational level (**Figure 1B**, cluster #4). We termed these 21 genes (listed in **Figure 6C**) ’effector’ genes and suggest that given the large pleiotropic effect excreted by the mutated eIF2B upon cytokine treatment, they may represent the gene set directly affected by the reduced regulatory capacity of Mut astrocytes. For most effector genes, intrinsically lower IS values in Mut relative to WT astrocytes (IS_WT0hr_ vs IS_Mut0hr_) combined with the stronger response of WT to cytokine activation (**Figure 6B**, p=0.03), synergistically resulted in a striking difference upon 24 hr of cytokine exposure (median IS_WT24hr_/IS_Mut24hr_ ∼8, **Figure 6B**, p=0.0004). Thus, collectively, by intersecting hundreds of genes affected by the mutated eIF2B with IS-specific alterations, we suggest a much shorter list of genes possibly directly compromised in Mut astrocytes. As would have been expected for drivers of the pervasive dysregulation observed, the effector genes are critically implicated in fundamental cell functions, including RNA-, protein-, lipid-, and energy-metabolism, vesicular trafficking, redox maintenance, cell cycle, cell-cell signaling, and cell death (**Figure 6C**). Notably, the effector genes are involved in neuronal development and plasticity, neural immuno-modulation, blood-brain-barrier maintenance, myelin sheath maintenance, and neuropathology (**Figure 6C**). Together, the data supports a working hypothesis, to be tested by future experiments, that a combined subtle dysregulation of some or all of the ’effector’ genes leads to VWMD pathology.

**Figure 4.**
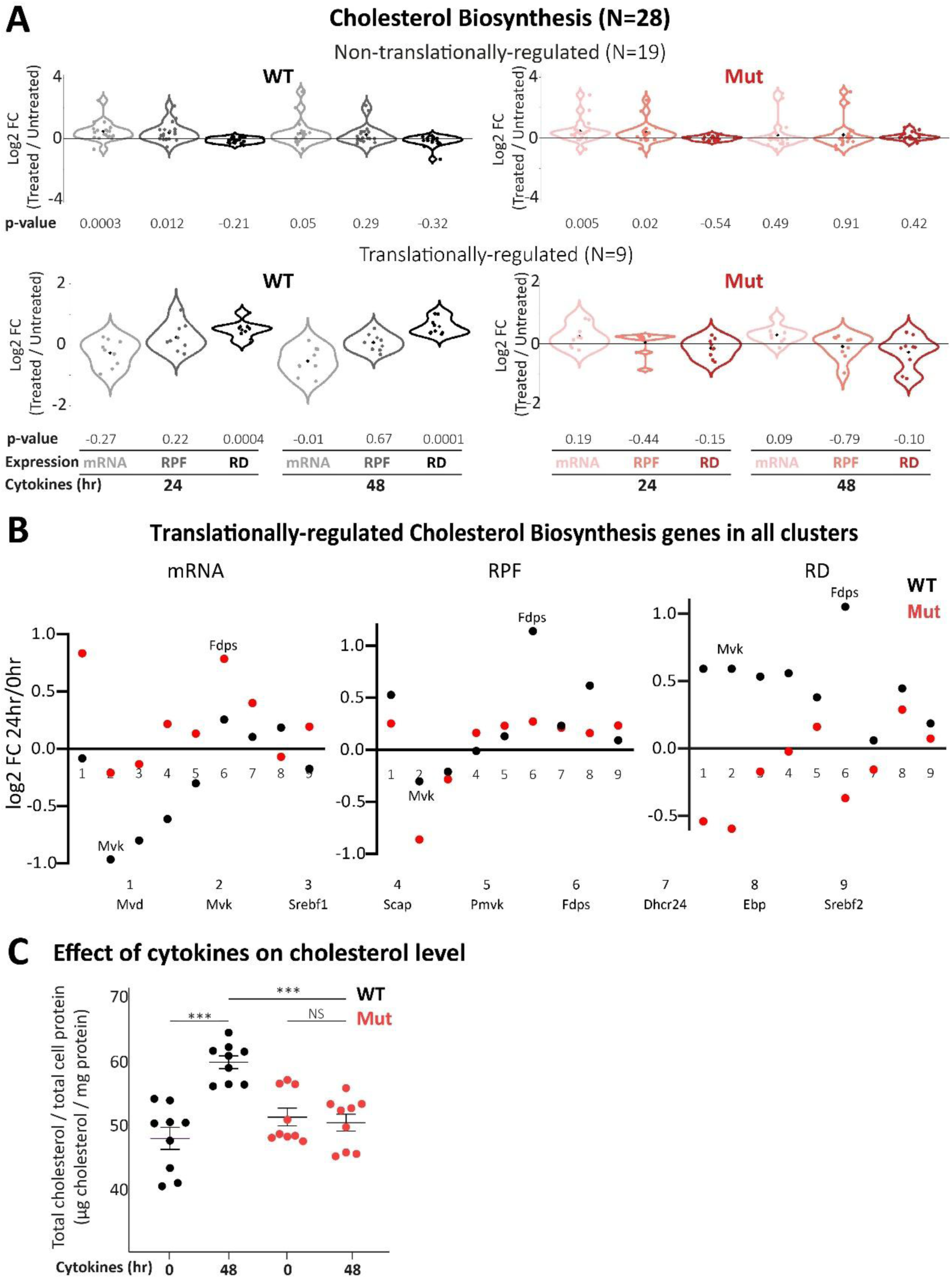
Effect of eIF2B mutation on expression response of genes related to cholesterol biosynthesis and its functional consequence. **(A)** Genes related to the cholesterol biosynthesis pathway (see Methods, N=28) classified based on their Log2FC RD value in WT at 48 hr were parted into two categories: (i) non-translationally regulated (RD<0.3; N=19, top) and (ii) translationally-regulated (RD>0.3; N=9; bottom). Log2 FC values obtained for WT and Mut are shown as detailed in Figure 1(C-E). **(B)** Log2 FC values of mRNA, RPF, and RD of each of the 9 translationally-regulated genes regardless of their cluster’s residency (Figure 1B), in response to 24 hr of cytokine treatment. (C) WT (black) and Mut (red) astrocytes were treated or not with cytokines for 48 hr followed by measurement of total cholesterol quantification (see Methods). Total cholesterol was normalized per total cells protein. P-values were determined by one-way ANOVA using Sidak correction for multiple comparisons. ***P ≤ 0.0002.

**Figure 5.**
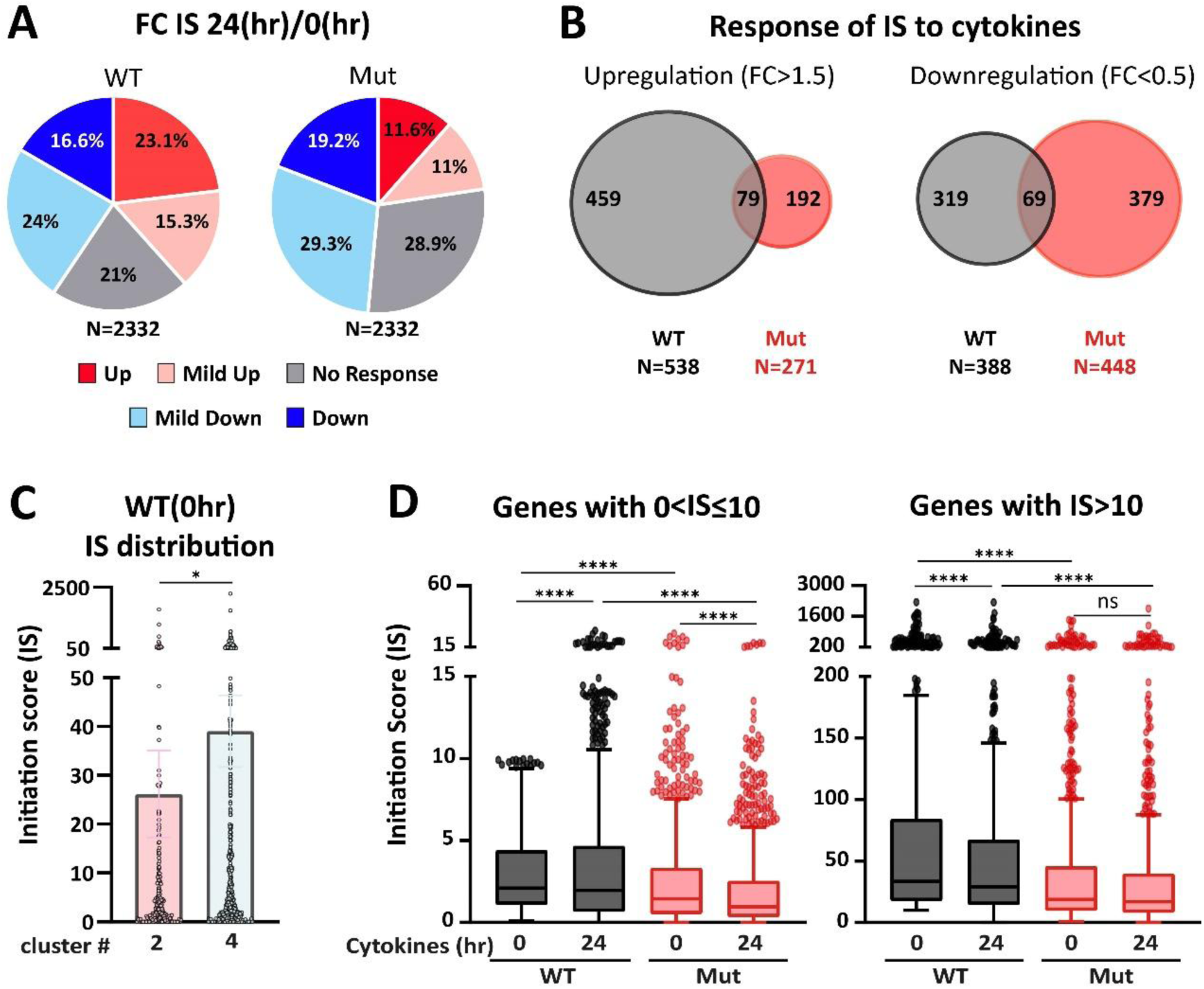
**(A)** Response of initiation score (IS) of 2332 genes to 24 hr of cytokine treatment in WT and Mut astrocytes. The FC IS(24hr)/IS(0hr) is presented in 5 categories: Upregulation (FC>1.5, dark red), mild up (1.2<FC≤1.5, light red), no response (0.8≤FC≤1.2, grey), mild down (0.5≤FC<0.8, light blue), and downregulated (FC<0.5, dark blue) **(B)** Venn diagram of the upregulated (FC>1.5) and downregulated (FC<0.5) IS values of genes in WT and Mut astrocytes, emphasizing the shared and unique responses. **(C)** IS distribution of WT(0hr) genes within clusters #2 and #4 (Fig. 1B). Kolmogorov-Smirnov test, *P=0.01 **(D)** IS distribution of genes whose IS in WT(0hr) is 0<IS≤10 (left) and IS >10 (right), within the four indicated datasets. 2-way Anova, ****P<0.0001.

**Figure 6.**
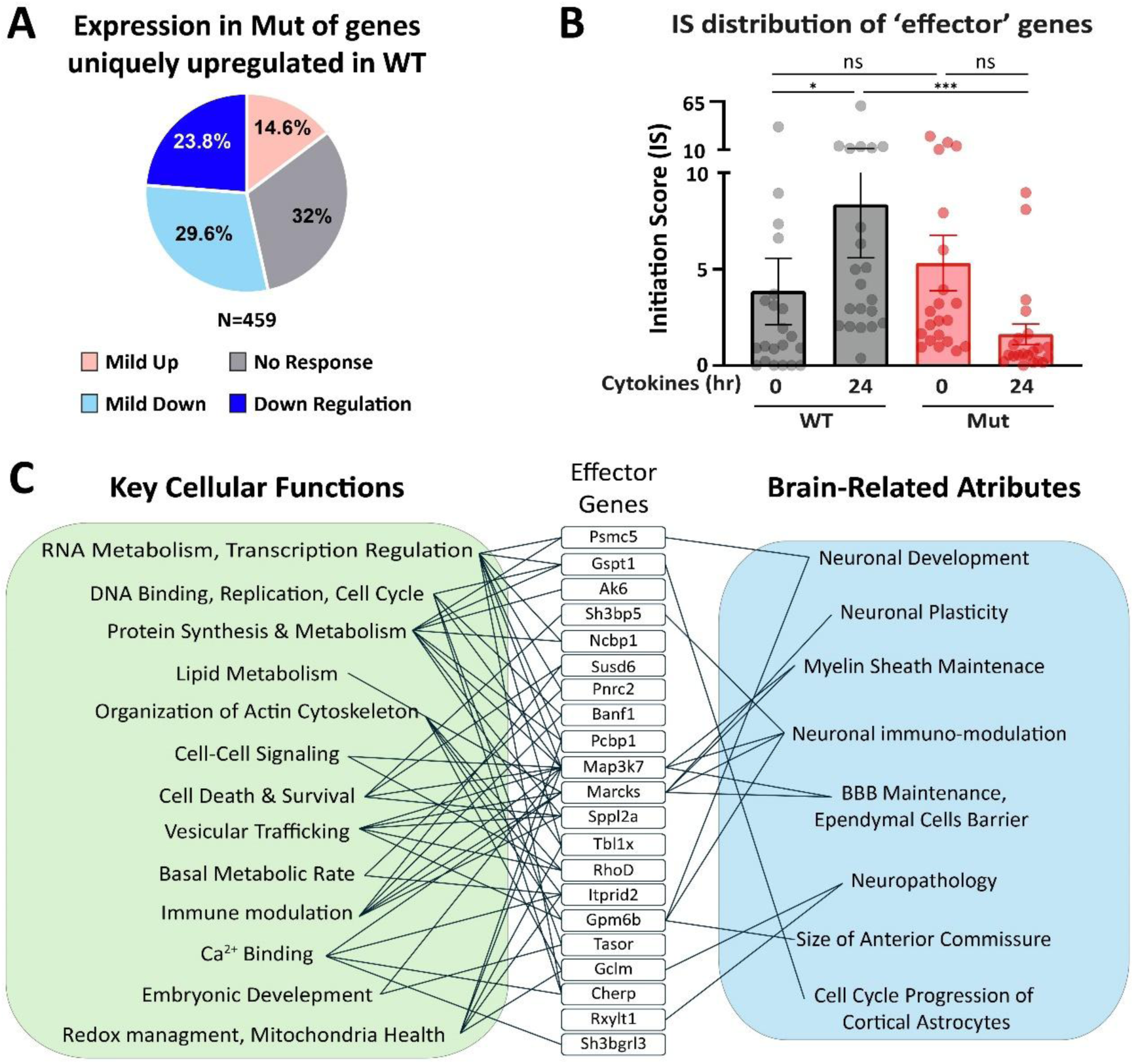
**(A)** FC IS Mut(24hr)/Mut(0hr) of the 459 genes uniquely upregulated (FC>1.5) in WT (Fig. 5B, left), divided into four categories: downregulated (FC<0.5, dark blue), mild down (0.5≤FC<0.8, light blue), no response (0.8≤FC<1.2, grey) and mild up (1.2≤FC<1.5, light red). **(B)** The distribution of the 21 ’Effector’ genes’ IS values within the four indicated datasets is shown. *P=0.03, ***P=0.0004, ns= not significant. **(C)** Key cellular functions and specific neurological attributes associated with the top dysregulated genes.

### Regulation of uORFs is also affected by eIF2B mutation and indirectly impacts the expression of main ORFs

While for all 21 ’effector’ gene candidates the cytokine-induced IS increase was compromised in Mut astrocytes, for seven of them, baseline IS in Mut was somewhat higher than in WT astrocytes (IS_Mut0hr_>IS_WT0hr_), i.e., regulation of their translation initiation is predicted to be more potent in Mut than WT before cytokine treatment. Interestingly, RPF distribution along these transcripts (Susd6, Banf1, Tbl1X, Gpm6b, TasoR, Pcbp1, and Ak6) demonstrates ribosomes occupancy upstream of the main start codon, suggesting the presence of regulatory uORFs that are dysregulated in Mut astrocytes. Given the broad implications of uORFs within the 5’UTRs for regulating main ORFs’ translation initiation, we zoomed out to test whether the anomaly affecting Mut’s translation initiation at the main ORF similarly affects the translation initiation of uORFs. To this end we used the trips-viz tool (15) to quantify uORFs’ translation. Our analysis (see Methods) revealed translation of >70K uORFs in our datasets, with top-ranked 5000 uORFs, which we further analyzed. The analysis showed that 33% and 30% of the uORFs start with an AUG codon in WT and Mut astrocytes, respectively (p<0.005; Chi-Square test). In support of the latter, inspection of the RPF distribution along the ATF4 5’UTR shows that initiation at the UUG of ORF1, known as non-regulatory (30), intensifies upon cytokine treatment more in Mut than in WT astrocytes. However, WT initiates more frequently at the downstream uORF2 and uORF3 than Mut (**Supplementary Figure S6**). Interestingly, we found that the genes exhibiting a translational response to cytokine activation (**Figure 1B**, cluster #4) are the most enriched with uORFs. 28% of cluster #4 genes harbor at least one uORF, compared to 26%, 23%, 25%, 21%, and 21% of the genes in clusters #1,2,3,5, and 6, respectively. Moreover, cluster #4 genes share differential characteristics of their 5’UTR, CDS, and 3’UTR relative to all others. Specifically, they harbor (i) higher GC content in the 5’UTR (66% vs. 64%; p < 1e-7); (ii) shorter CDS (average of 1523 nt vs. 1962 nt; p < 1e-30), and (iii) longer 3’UTR (1647 nt vs. 1360 nt; p < 1e-15). Future dedicated experiments will identify yet-unknown *cis*-regulatory elements that confer sensitivity to eIF2B function.

## Discussion

This study constitutes the first translatome-based infrastructure for discovering the molecular mechanisms involved in VWMD pathology. In addition, it provides a profound insight into the importance of eIF2B as a mediator of gene-specific translation regulation during cytokine-mediated activation of astrocytes, a critical biological process. The first take-home message is that even a slight partial loss of eIF2B function due to homozygosity of the mild eIF2B5(R132H) mutation (11) is sufficient to effectively disrupt translation regulation programs required for cytokine-mediated activation of astrocytes, leading to functional deficits. A more general insight is the upregulation of mRNA abundance as a common compensatory strategy upon a failure to enhance the translation of specific gene products. The effect of eIF2B5(R132H) mutation on the transcriptome’s composition was initially detected over a decade ago using expression microarray analyses of whole WT and Mut brains at different postnatal developmental stages (31). Both studies substantiate the conclusion that transcriptome changes caused by the malfunction of a translation initiation factor denote the involvement of indirect effects on RNA metabolism. These must be mediated by the dysregulated translation of a small group of effector genes, which affect expression networks and biochemical pathways of downstream genes. Several scenarios are possible, such as sub-optimal levels of RNA binding proteins involved in mechanisms governing co-translational mRNA decay (32–34) combined with compensatory responses aiming at optimal homeostasis. An example is the metabolic change acquired by Mut astrocytes owing to their hypervulnerability to energy deficit and oxidative stress (21).

An important insight relates to the sensitivity of RP and OXPHOS genes to eIF2B mutation. RPs, the building blocks of ribosomes, are generated in response to demand for increasing the production of necessary proteins, as in the case of cytokine-mediated astrocyte activation. Since mRNA translation is energetically expensive, a parallel increase in mitochondrial oxidative phosphorylation is required to upsurge ATP levels. Therefore, it is unsurprising that both gene groups are translationally upregulated in response to acute demand mediated by the activation of the mTOR axis (10,35). mTORC1 supports global protein synthesis by increasing the availability of translation initiation eIF4E, the 5’cap-binding protein (1,36), generally accepted as the rate-limiting step of translation initiation under normal conditions. The transcripts of both RP and OXPHOS gene groups are equipped with *cis*-regulatory elements, which render them highly dependent on the availability of eIF4E. For RPs it is the 5’-terminal oligopyrimidines tract (5’TOP) (19,23,37), and for OXPHOS it is the Translation Initiator of Short 5′UTR (TISU) element (38–40). The current study shows for the first time that the hypomorphic eIF2B mutation renders TC availability the rate-limiting factor for the upregulation of the RPs and OXPHOS gene subsets in response to cytokine-mediated activation. Interestingly, this phenomenon emphasizes the understanding that maximal eIF2B enzymatic activity in WT astrocytes is required to support appropriate astrocytic activation.

The study shows that even a mild loss-of-function mutation in eIF2B negatively affects the translation of transcripts, which are typically subjected to the most significant rise upon acute demands. Yet, the data indicates a broad sensitivity range of members within the RP and OXPHOS functional gene sets (**Figures 2AB and 3AB**), implying that an insufficient number of functional ribosomes (**Figure 2D**) and ETC (**Figure 3D**) can arise from dysregulated translation of only a few members of these multi-subunit complexes. The Cox6c case (Figure 3C) supports the notion that an accurate translation rate via uORF-mediated attenuation is essential for generating multi-subunit complexes. Interestingly, 34% of the OXPHOS genes within the translationally regulated group harbor uORFs compared to 11.5% of the OXPHOS genes in the non-translationally regulated group (Chi-square p-value = 0.036) (Figure 3A), advocating a differential effect of eIF2B mutation. The cholesterol biosynthesis pathway referred to in the current study provides a proof-of-concept for a similar phenomenon related to multi-component biochemical pathways, as it shows that a mild dysregulation of only a few enzymes in this pathway leads to a detrimental effect (**Figure 4**).

Cholesterol is a key structural element of myelin sheaths, which exhibit a defective phenotype in VWMD. Although the blood-brain-barrier essentially prevents the influx of cholesterol from the bloodstream, the adult brain is the most cholesterol-rich organ in the body (41), mainly owing to cholesterol biosynthesis in astrocytes which provide it to neighboring cells upon demand (42). The significance of cholesterol biosynthesis in astrocytes is crucial for re-myelination (43). In addition to its being an essential component of cellular membranes, cholesterol within the brain also serves as a precursor for various molecules such as steroid hormones. Notably, the energy demand for cholesterol biosynthesis is very high, requiring the utilization of 18 acetyl-CoA molecules, 18 ATP molecules, and 25 reduction equivalents (NADPH/NADH) per single cholesterol molecule (44). Cholesterol homeostasis is tightly regulated at the transcriptional and post-transcriptional levels (29), while information related to regulation at the translation level is limited. Our novel finding that translation of 9 of 28 genes of the cholesterol biosynthesis pathway exhibit sensitivity to the eIF2B mutation (Figure 4AB) strongly suggests it plays a role in VWMD pathology since cholesterol biosynthesis in astrocytes is connected to the process of white matter regeneration in the brain (28). Defective myelin maintenance is echoed by slow and defective myelination and re-myelination in eIF2B5^R132H/R132H^ mice (11,25). The inability of Mut astrocytes to synthesize enough cholesterol upon demand (**Figure 4C**) also agrees with the defective morphology and blunt processes of Mut astrocytes and oligodendrocytes (45,46) (47).

Many features may influence the sensitivity of transcripts to TC levels generated by eIF2B. A likely *cis*-acting factor is the primary sequence around the mRNA’s start codon, which facilitates its base-pairing with the rRNA to stabilize the start codon position at the P-site of the ribosome and consequent codon-anticodon base pairing to promote an effective initiation event. The IS of each ORF represents a specific strength of this feature. Additional *cis-*acting features may affect the combined sensitivity to all potential changes in the availability of *trans*-acting factors, which include the TC level, among others (48). The *trans*-acting factors may directly affect the initiation at main ORFs (as in the case of RPL19; **Figure 2C**) or indirectly via initiating at the start codon of regulatory uORFs (as in the case of Cox6c; **Figure 3C**). The Cox6c uORF possibly functions to control the rate of its synthesis tightly. The dysregulated uORF-mediated translation attenuation of Cox6c synthesis in Mut may negatively affect its balance with other subunits of the ETC IV complex, negatively affecting its assembly, integration into the mitochondria inner membrane, and consequent enzymatic activity. Since ISR-regulating uORFs are included in the possibilities mentioned above, hyperactivation of the ISR response by Mut astrocytes is expected. However, this is not always the case, as the current study shows that cytokine-mediated activation elicits stimulation of the classical ISR markers (22) in both WT and Mut astrocytes without further hyperactivation in Mut (**Figure 1C**). A hyperactive ISR is generally perceived as the ultimate reason for VWMD pathology (49–52). However, this statement has some inconsistent indications (53). The current study shows that the molecular consequences leading to VWMD, at least in response to scenarios involving the activation of astrocytes by pro-inflammatory cytokines, which mimics what occurs in the brain in response to multiple physiological cues (10,54,55), must include scenarios beyond the hyperactivation of ISR. Moreover, The data positions some hypomorphic eIF2B mutations-driven dysregulation in VWMD, as an upstream driver of manifested ISR.

We show here that in addition to the hyper-sensitivity to stress, a simple lack of ability to increase the translation of specific transcripts in response to acute demand is an essential component of VWMD. While initial evidence supporting this concept was provided over a decade ago (45), the current study offers a supportive meta-genomic analysis in codon resolution. Future extensive experimental work for solving such a complex mechanism should include (i) dissection of the exact 5’UTR features assigning high sensitivity to mild changes in TC levels; and (ii) investigation of physiological scenarios that involve changes in eIF2B activity in the absence of ISR, such as via GSK3- or CK2-mediated direct phosphorylation of eIF2B subunits (56), or via binding of natural ligands such as sugar phosphates (57).

Along these lines, in addition to some members of RP, OXPHOS, and cholesterol biosynthesis gene sets, the current study suggests the involvement of 21 specific ’effector’ genes whose translation stimulation in astrocytes is severely lacking in response to cytokine-mediated activation (**Figure 6C**). A key ’effector’ example is myristoylated alanine-rich C-kinase substrate (Marcks), a PKC target. Marcks is involved in: (i) transduction of cytokine and other neuro-immune modulatory signals mediated by PKC in astrocytes (58); (ii) regulation of myelin sheath formation in response to extracellular stimuli (59); (iii) ability of Ependymal cells to maintain an effective cerebrospinal fluid-brain barrier (60); (iv) development of synaptic spines (61) and regrowth of catecholaminergic and serotonergic axons (62). Another key ’effector’ example is Gclm, the first rate-limiting enzyme of glutathione synthesis. Glutathione (GSH) is an essential antioxidant for protection from reactive oxygen species (ROS). GSH production, utilization, and export balance become critical during cytokine exposure. Dysregulation of Gclm translation can severaly compromise astrocytic support functions and overall brain resilience to stress and injury. The latter agrees with the hypersensitivity of eIF2B-Mut astrocytes to oxidative stress (21).

It is important to note that we used cultured astrocytes isolated from newborn mice, while astrocytes in the adult brain are highly heterogeneous and often translate mRNAs in specific sub-cellular compartments to ensure their function and polarity (63,64). However, it is tempting to speculate that most of the significant anomalies observed in this study are likely to occur also in the brains of human VWMD patients. Future experiments will reveal the links of ’effector’ gene candidates to VWMD pathology and provide opportunities for intervention by drug repurposing and developing new therapeutic modalities.

## Supporting information

Mandelboum et al - Supplementary Figures

## Acknowledgements

We thank Ran Elkon for computation backing, Irit Gat-Viks for valuable discussions, Oded Meyuhas for thoughtful conversations and critical reading of the manuscript, and VWMD families for their support.

## Abbreviations

5’UTR: 5’-untranslated region
CDS: coding sequence
ETC: electron transfer chain
IS: initiation score
ISR: integrated stress response
OXPHOS: oxidative phosphorylation
RD: ribosome density
RP: ribosomal proteins
RRO: relative ribosome occupancy
TC: eIF2·GTP·Met-tRNA^Met^ ternary complex
uAUG: upstream AUG
uORF: upstream open reading frame
VWMD: vanishing white matter disease.

## Supplementary Data

- A pdf file containing Supplementary Figures S1-S7.

- Supplementary Table S1 (SuperSeries GSE216583; WT SubSeries GSE216582; Mut Subseries GSE252362). Excel file containing RNA-seq and Ribo-seq data of WT and eIF2B5-R132H-R132H mutant astrocytes, with an overall coverage of 12,205 genes. For each gene, the Ensemble ID and symbol are specified. The first tab of the Excel file presents the log2 fold change value of each mRNA and RPF upon (24-hr-treated vs untreated), and (48-hr-treated vs untreated) together with the TE value of each gene. The 2nd-7th tabs of the Excel file present specific gene lists that were differentially expressed within the tested samples. The 8th tab of the Excel file presents specific P-values of subsets of genes compared with the expression of all the rest of the genes in the dataset. P-values were calculated using Wilcoxon’s test.

## References

1. Saxton, R.A. and Sabatini, D.M. (2017) mTOR Signaling in Growth, Metabolism, and Disease. Cell, 169, 361–371.

2. Pavitt, G.D. (2018) Regulation of translation initiation factor eIF2B at the hub of the integrated stress response. Wiley Interdiscip Rev RNA, 9, e1491.

3. Marintchev, A. and Ito, T. (2020) eIF2B and the Integrated Stress Response: A Structural and Mechanistic View. Biochemistry, 59, 1299–1308.

4. Baird, T.D. and Wek, R.C. (2012) Eukaryotic initiation factor 2 phosphorylation and translational control in metabolism. Adv Nutr, 3, 307–321.

5. Leegwater, P.A., Vermeulen, G., Konst, A.A., Naidu, S., Mulders, J., Visser, A., Kersbergen, P., Mobach, D., Fonds, D., van Berkel, C.G., et al. (2001) Subunits of the translation initiation factor eIF2B are mutant in leukoencephalopathy with vanishing white matter. Nat Genet, 29, 383–388.

6. Hamilton, E.M.C., van der Lei, H.D.W., Vermeulen, G., Gerver, J.A.M., Lourenco, C.M., Naidu, S., Mierzewska, H., Gemke, R., de Vet, H.C.W., Uitdehaag, B.M.J., et al. (2018) Natural History of Vanishing White Matter. Ann Neurol, 84, 274–288.

7. Dooves, S., Bugiani, M., Postma, N.L., Polder, E., Land, N., Horan, S.T., van Deijk, A.L., van de Kreeke, A., Jacobs, G., Vuong, C., et al. (2016) Astrocytes are central in the pathomechanisms of vanishing white matter. J Clin Invest, 126, 1512–1524.

8. Bugiani, M., Vuong, C., Breur, M. and van der Knaap, M.S. (2018) Vanishing white matter: a leukodystrophy due to astrocytic dysfunction. Brain Pathol, 28, 408–421.

9. Man, J.H.K., van Gelder, C., Breur, M., Okkes, D., Molenaar, D., van der Sluis, S., Abbink, T., Altelaar, M., van der Knaap, M.S. and Bugiani, M. (2022) Cortical Pathology in Vanishing White Matter. Cells, 11.

10. Mandelboum, S., Herrero, M., Atzmon, A., Ehrlich, M. and Elroy-Stein, O. (2023) Effective extraction of polyribosomes exposes gene expression strategies in primary astrocytes. Nucleic Acids Res, 51, 3375–3390.

11. Geva, M., Cabilly, Y., Assaf, Y., Mindroul, N., Marom, L., Raini, G., Pinchasi, D. and Elroy-Stein, O. (2010) A mouse model for eukaryotic translation initiation factor 2B-leucodystrophy reveals abnormal development of brain white matter. Brain, 133, 2448–2461.

12. McGlincy, N.J. and Ingolia, N.T. (2017) Transcriptome-wide measurement of translation by ribosome profiling. Methods, 126, 112–129.

13. Kim, D., Pertea, G., Trapnell, C., Pimentel, H., Kelley, R. and Salzberg, S.L. (2013) TopHat2: accurate alignment of transcriptomes in the presence of insertions, deletions and gene fusions. Genome Biol, 14, R36.

14. Liao, Y., Smyth, G.K. and Shi, W. (2014) featureCounts: an efficient general purpose program for assigning sequence reads to genomic features. Bioinformatics, 30, 923–930.

15. Kiniry, S.J., O’Connor, P.B.F., Michel, A.M. and Baranov, P.V. (2019) Trips-Viz: a transcriptome browser for exploring Ribo-Seq data. Nucleic Acids Res, 47, D847–D852.

16. Mandelboum, S., Manber, Z., Elroy-Stein, O. and Elkon, R. (2019) Recurrent functional misinterpretation of RNA-seq data caused by sample-specific gene length bias. PLoS Biol, 17, e3000481.

17. Chothani, S., Adami, E., Ouyang, J.F., Viswanathan, S., Hubner, N., Cook, S.A., Schafer, S. and Rackham, O.J.L. (2019) deltaTE: Detection of Translationally Regulated Genes by Integrative Analysis of Ribo-seq and RNA-seq Data. Curr Protoc Mol Biol, 129, e108.

18. Liebermeister, W., Noor, E., Flamholz, A., Davidi, D., Bernhardt, J. and Milo, R. (2014) Visual account of protein investment in cellular functions. Proc Natl Acad Sci U S A, 111, 8488–8493.

19. Perry, R.P. (2005) The architecture of mammalian ribosomal protein promoters. BMC Evol Biol, 5, 15.

20. Choudhary, S., Li, W. and A, D.S. (2020) Accurate detection of short and long active ORFs using Ribo-seq data. Bioinformatics, 36, 2053–2059.

21. Herrero, M., Daw, M., Atzmon, A. and Elroy-Stein, O. (2021) The Energy Status of Astrocytes Is the Achilles’ Heel of eIF2B-Leukodystrophy. Cells, 10.

22. Sims, S.G., Cisney, R.N., Lipscomb, M.M. and Meares, G.P. (2022) The role of endoplasmic reticulum stress in astrocytes. Glia, 70, 5–19.

23. Meyuhas, O. and Kahan, T. (2015) The race to decipher the top secrets of TOP mRNAs. Biochim Biophys Acta, 1849, 801–811.

24. Buccitelli, C. and Selbach, M. (2020) mRNAs, proteins and the emerging principles of gene expression control. Nat Rev Genet, 21, 630–644.

25. Gat-Viks, I., Geiger, T., Barbi, M., Raini, G. and Elroy-Stein, O. (2015) Proteomics-level analysis of myelin formation and regeneration in a mouse model for Vanishing White Matter disease. J Neurochem, 134, 513–526.

26. Raini, G., Sharet, R., Herrero, M., Atzmon, A., Shenoy, A., Geiger, T. and Elroy-Stein, O. (2017) Mutant eIF2B leads to impaired mitochondrial oxidative phosphorylation in vanishing white matter disease. J Neurochem, 141, 694–707.

27. Elroy-Stein, O. and Schiffmann, R. (2020) In Rosenberg, R. N. and Pascual, J. M. (eds.), Rosenberg’s Molecular and Genetic Basis of Neurological and Psychiatric Diseases. 6th ed. Elsevier Academic Press Inc., Cambridge, MA, USA, Vol. 2, pp. 301-317.

28. Berghoff, S.A., Spieth, L. and Saher, G. (2022) Local cholesterol metabolism orchestrates remyelination. Trends Neurosci, 45, 272–283.

29. Sharpe, L.J. and Brown, A.J. (2013) Controlling cholesterol synthesis beyond 3-hydroxy-3-methylglutaryl-CoA reductase (HMGCR). J Biol Chem, 288, 18707–18715.

30. Zhang, F. and Hinnebusch, A.G. (2011) An upstream ORF with non-AUG start codon is translated in vivo but dispensable for translational control of GCN4 mRNA. Nucleic Acids Res, 39, 3128–3140.

31. Marom, L., Ulitsky, I., Cabilly, Y., Shamir, R. and Elroy-Stein, O. (2011) A point mutation in translation initiation factor eIF2B leads to function--and time-specific changes in brain gene expression. PLoS One, 6, e26992.

32. Pelechano, V., Wei, W. and Steinmetz, L.M. (2015) Widespread Co-translational RNA Decay Reveals Ribosome Dynamics. Cell, 161, 1400–1412.

33. Gentilella, A., Moron-Duran, F.D., Fuentes, P., Zweig-Rocha, G., Riano-Canalias, F., Pelletier, J., Ruiz, M., Turon, G., Castano, J., Tauler, A. et al. (2017) Autogenous Control of 5’TOP mRNA Stability by 40S Ribosomes. Mol Cell, 67, 55–70 e54.

34. Hopfler, M. and Hegde, R.S. (2023) Control of mRNA fate by its encoded nascent polypeptide. Mol Cell, 83, 2840–2855.

35. Latacz, A., Russell, J.A., Oclon, E., Zubel-lojek, J. and Pierzchala-Koziec, K. (2015) mTOR Pathway - Novel Modulator of Astrocyte Activity. Folia Biol (Krakow), 63, 95–105.

36. Iadevaia, V., Liu, R. and Proud, C.G. (2014) mTORC1 signaling controls multiple steps in ribosome biogenesis. Semin Cell Dev Biol, 36, 113–120.

37. Philippe, L., Vasseur, J.J., Debart, F. and Thoreen, C.C. (2018) La-related protein 1 (LARP1) repression of TOP mRNA translation is mediated through its cap-binding domain and controlled by an adjacent regulatory region. Nucleic Acids Res, 46, 1457–1469.

38. Dikstein, R. (2012) Transcription and translation in a package deal: the TISU paradigm. Gene, 491, 1–4.

39. Haimov, O., Sinvani, H., Martin, F., Ulitsky, I., Emmanuel, R., Tamarkin-Ben-Harush, A., Vardy, A. and Dikstein, R. (2017) Efficient and Accurate Translation Initiation Directed by TISU Involves RPS3 and RPS10e Binding and Differential Eukaryotic Initiation Factor 1A Regulation. Mol Cell Biol, 37.

40. Havkin-Solomon, T., Fraticelli, D., Bahat, A., Hayat, D., Reuven, N., Shaul, Y. and Dikstein, R. (2023) Translation regulation of specific mRNAs by RPS26 C-terminal RNA-binding tail integrates energy metabolism and AMPK-mTOR signaling. Nucleic Acids Res, 51, 4415–4428.

41. Dietschy, J.M. (2009) Central nervous system: cholesterol turnover, brain development and neurodegeneration. Biol Chem, 390, 287–293.

42. Orth, M. and Bellosta, S. (2012) Cholesterol: its regulation and role in central nervous system disorders. Cholesterol, 2012, 292598.

43. Molina-Gonzalez, I., Holloway, R.K., Jiwaji, Z., Dando, O., Kent, S.A., Emelianova, K., Lloyd, A.F., Forbes, L.H., Mahmood, A., Skripuletz, T. et al. (2023) Astrocyte-oligodendrocyte interaction regulates central nervous system regeneration. Nat Commun, 14, 3372.

44. Saher, G., Quintes, S. and Nave, K.A. (2011) Cholesterol: a novel regulatory role in myelin formation. Neuroscientist, 17, 79–93.

45. Cabilly, Y., Barbi, M., Geva, M., Marom, L., Chetrit, D., Ehrlich, M. and Elroy-Stein, O. (2012) Poor cerebral inflammatory response in eIF2B knock-in mice: implications for the aetiology of vanishing white matter disease. PLoS One, 7, e46715.

46. Bugiani, M., Boor, I., van Kollenburg, B., Postma, N., Polder, E., van Berkel, C., van Kesteren, R.E., Windrem, M.S., Hol, E.M., Scheper, G.C., et al. (2011) Defective glial maturation in vanishing white matter disease. J Neuropathol Exp Neurol, 70, 69–82.

47. Herrero, M., Mandelboum, S. and Elroy-Stein, O. (2019) eIF2B Mutations Cause Mitochondrial Malfunction in Oligodendrocytes. Neuromolecular Med, 21, 303–313.

48. Kim, Y.S., Kimball, S.R., Piskounova, E., Begley, T.J. and Hempel, N. (2024) Stress response regulation of mRNA translation: Implications for antioxidant enzyme expression in cancer. Proc Natl Acad Sci U S A, 121, e2317846121.

49. Abbink, T.E.M., Wisse, L.E., Jaku, E., Thiecke, M.J., Voltolini-Gonzalez, D., Fritsen, H., Bobeldijk, S., Ter Braak, T.J., Polder, E., Postma, N.L. et al. (2019) Vanishing white matter: deregulated integrated stress response as therapy target. Ann Clin Transl Neurol, 6, 1407–1422.

50. van der Voorn, J.P., van Kollenburg, B., Bertrand, G., Van Haren, K., Scheper, G.C., Powers, J.M. and van der Knaap, M.S. (2005) The unfolded protein response in vanishing white matter disease. J Neuropathol Exp Neurol, 64, 770–775.

51. van Kollenburg, B., van Dijk, J., Garbern, J., Thomas, A.A., Scheper, G.C., Powers, J.M. and van der Knaap, M.S. (2006) Glia-specific activation of all pathways of the unfolded protein response in vanishing white matter disease. J Neuropathol Exp Neurol, 65, 707–715.

52. Wong, Y.L., LeBon, L., Basso, A.M., Kohlhaas, K.L., Nikkel, A.L., Robb, H.M., Donnelly-Roberts, D.L., Prakash, J., Swensen, A.M., Rubinstein, N.D. et al. (2019) eIF2B activator prevents neurological defects caused by a chronic integrated stress response. Elife, 8.

53. Wisse, L.E., Ter Braak, T.J., van de Beek, M.C., van Berkel, C.G.M., Wortel, J., Heine, V.M., Proud, C.G., van der Knaap, M.S. and Abbink, T.E.M. (2018) Adult mouse eIF2Bepsilon Arg191His astrocytes display a normal integrated stress response in vitro. Sci Rep, 8, 3773.

54. Hyvarinen, T., Hagman, S., Ristola, M., Sukki, L., Veijula, K., Kreutzer, J., Kallio, P. and Narkilahti, S. (2019) Co-stimulation with IL-1beta and TNF-alpha induces an inflammatory reactive astrocyte phenotype with neurosupportive characteristics in a human pluripotent stem cell model system. Sci Rep, 9, 16944.

55. Giovannoni, F. and Quintana, F.J. (2020) The Role of Astrocytes in CNS Inflammation. Trends Immunol, 41, 805–819.

56. Wang, X., Paulin, F.E., Campbell, L.E., Gomez, E., O’Brien, K., Morrice, N. and Proud, C.G. (2001) Eukaryotic initiation factor 2B: identification of multiple phosphorylation sites in the epsilon-subunit and their functions in vivo. EMBO J, 20, 4349-4359.

57. Hao, Q., Heo, J.M., Nocek, B.P., Hicks, K.G., Stoll, V.S., Remarcik, C., Hackett, S., LeBon, L., Jain, R., Eaton, D. et al. (2021) Sugar phosphate activation of the stress sensor eIF2B. Nat Commun, 12, 3440.

58. Vitkovic, L., Aloyo, V.J., Maeda, S., Benzil, D.L., Bressler, J.P. and Hilt, D.C. (2005) Identification of myristoylated alanine-rich C kinase substrate (MARCKS) in astrocytes. Front Biosci, 10, 160–165.

59. 59. Siskova, Z., Baron, W., de Vries, H. and Hoekstra, D. (2006) Fibronectin impedes "myelin" sheet-directed flow in oligodendrocytes: a role for a beta 1 integrin-mediated PKC signaling pathway in vesicular trafficking. Mol Cell Neurosci, 33, 150–159.

60. Muthusamy, N., Sommerville, L.J., Moeser, A.J., Stumpo, D.J., Sannes, P., Adler, K., Blackshear, P.J., Weimer, J.M. and Ghashghaei, H.T. (2015) MARCKS-dependent mucin clearance and lipid metabolism in ependymal cells are required for maintenance of forebrain homeostasis during aging. Aging Cell, 14, 764–773.

61. Garrett, A.M., Schreiner, D., Lobas, M.A. and Weiner, J.A. (2012) gamma-protocadherins control cortical dendrite arborization by regulating the activity of a FAK/PKC/MARCKS signaling pathway. Neuron, 74, 269–276.

62. Theis, T., Kumar, S., Wei, E., Nguyen, J., Glynos, V., Paranjape, N., Askarifirouzjaei, H., Khajouienejad, L., Berthiaume, F., Young, W. and Schachner, M. (2020) Myristoylated alanine-rich C-kinase substrate effector domain peptide improves sex-specific recovery and axonal regrowth after spinal cord injury. FASEB J, 34, 12677–12690.

63. Cohen-Salmon, M., Slaoui, L., Mazare, N., Gilbert, A., Oudart, M., Alvear-Perez, R., Elorza-Vidal, X., Chever, O. and Boulay, A.C. (2021) Astrocytes in the regulation of cerebrovascular functions. Glia, 69, 817–841.

64. Sakers, K., Lake, A.M., Khazanchi, R., Ouwenga, R., Vasek, M.J., Dani, A. and Dougherty, J.D. (2017) Astrocytes locally translate transcripts in their peripheral processes. Proc Natl Acad Sci U S A, 114, E3830–E3838.

