## Supplementary material for "The Translational Landscape of Reactive Astrocytes Reveals the Impact of eIF2B-mediated Dysregulation in VWM Disease": Mandelboum et al - Supplementary Figures

### Figure S1

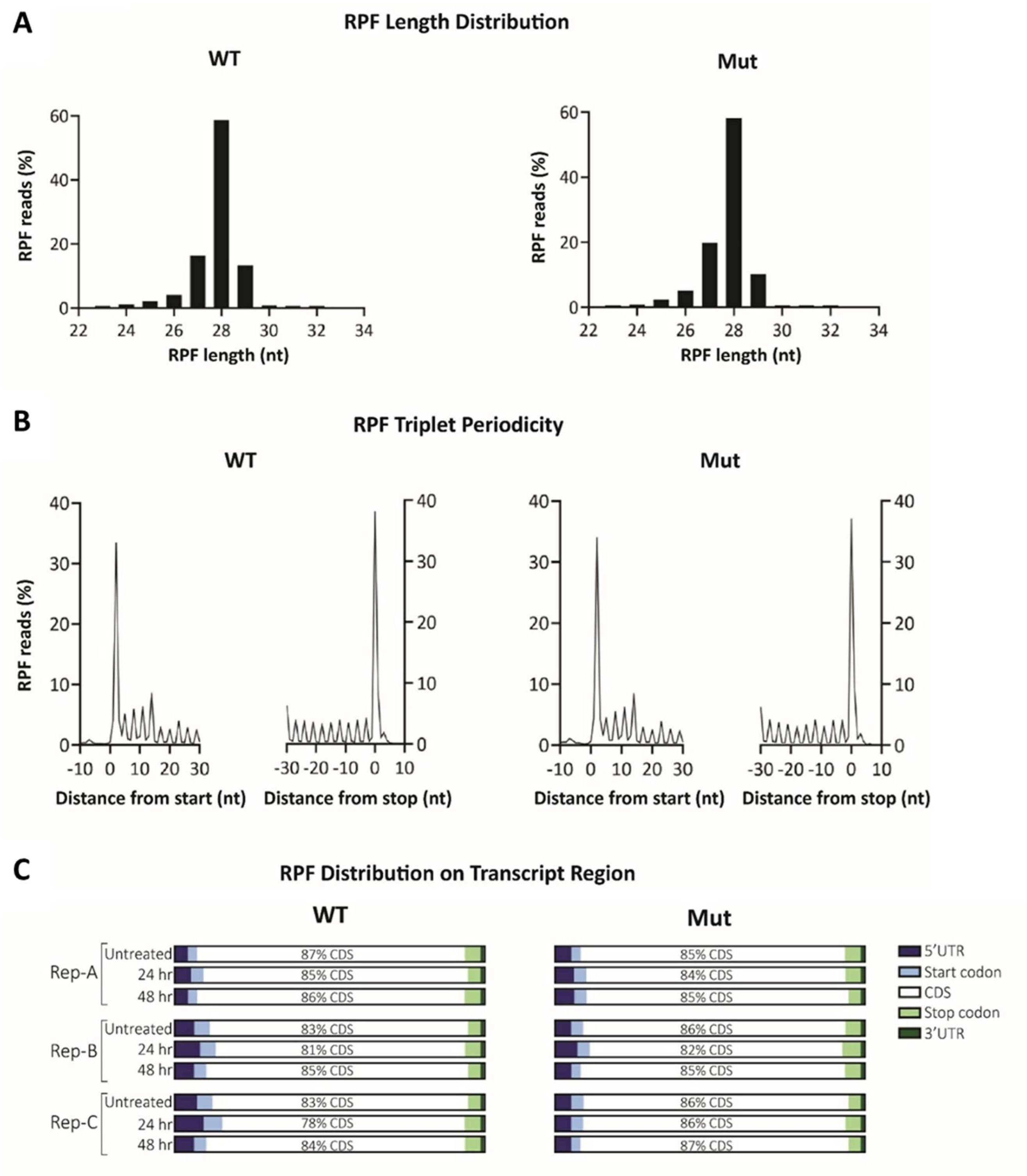


Figure S1

Quality control of Ribo-seq data. (A) Length distribution of RPF reads. 90% of the reads are 25-30 nt long, similar to the length of the mRNA region protected by a translating ribosome. (B) Triplet periodicity analysis of the protein coding transcripts. (C) Distribution of RPF reads along the different transcript regions. 5'UTR, 5' untranslated region; CDS, coding sequence; 3'UTR, 3'untrasnlated region. The data is presented for each replicate (Rep) independently.

Figure S2

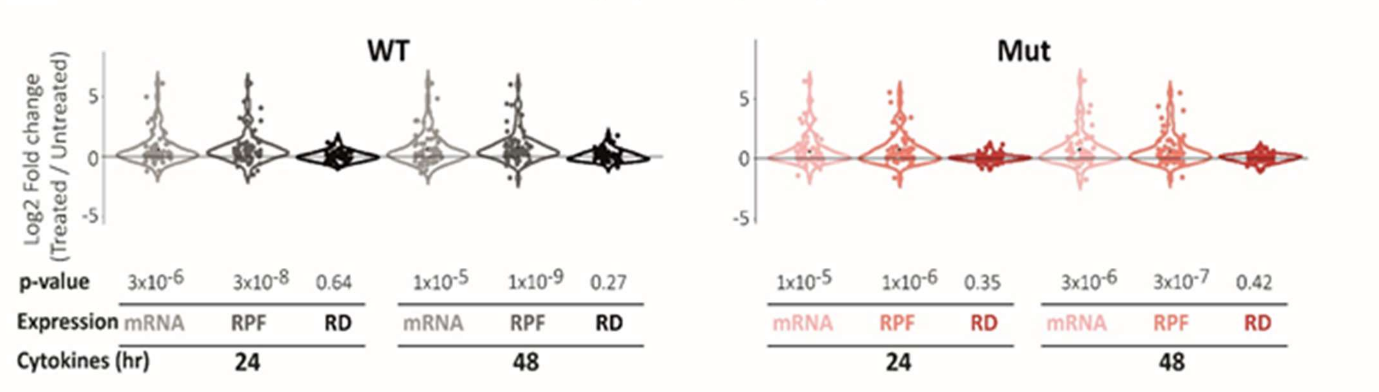


Figure S2

As detailed in Figure 1(C-E), shown are log2 fold-change values of total mRNA level, RPF, and RD of individual members of the 'KEGG Apoptosis' subset (N = 67), obtained in response to 24 and 48 hr cytokine treatment relative to untreated WT and eIF2B-mutated (Mut) astrocytes. P-values were calculated using Wilcoxon's test, comparing the selected gene sets to the rest of the genes in the dataset.

Figure S3

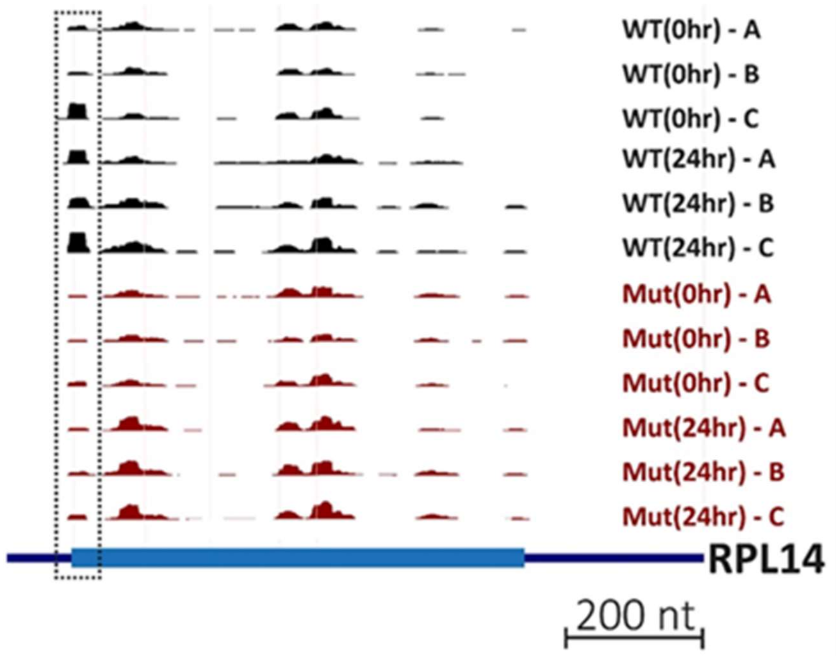


Figure S3

RPF map of RPL14, highlighting the increase in RPF reads at the AUG start codon, in WT but not in Mut, in response to 24 hr of cytokine treatment (dashed rectangle).

**Figure S4**

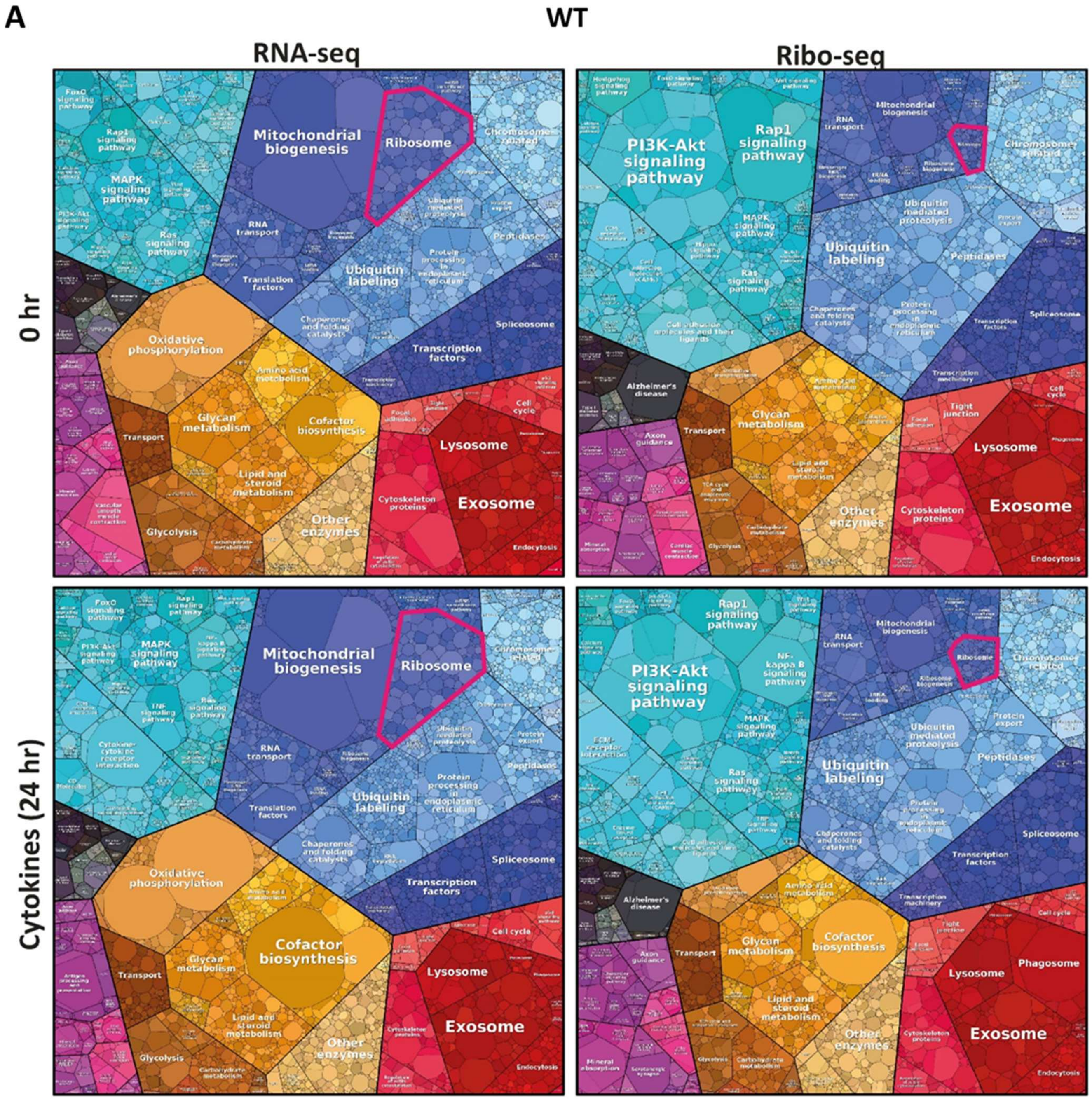
A
WT
RNA-seq
Ribo-seq
0 hr
Cytokines (24 hr)
Mitochondrial biogenesis
Ribosome
Ubiquitin labeling
Oxidative phosphorylation
Glycan metabolism
Cofactor biosynthesis
Lipid and steroid metabolism
Other enzymes
Lysosome
Exosome
Cytoskeleton proteins
Spliceosome
Transcription factors
Transport
Glycolysis
PI3K-Akt signaling pathway
Rap1 signaling pathway
MAPK signaling pathway
Ras signaling pathway
Chromosome related
Peptidases
Alzheimer's disease
Endocytosis
Phagosome
Cell cycle

B

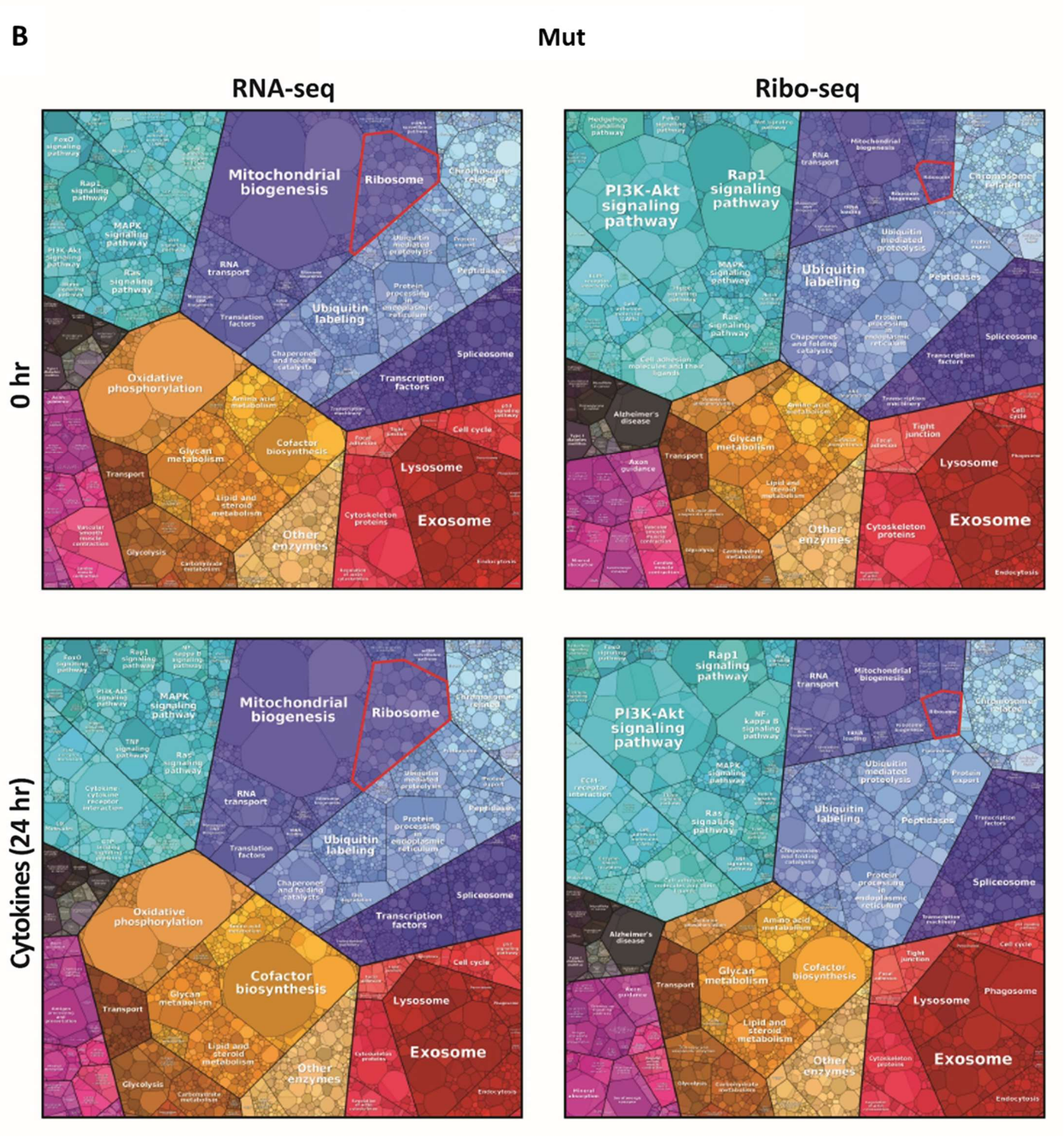


C

| | WT | | | Mut | | |
|---|---|---|---|---|---|---|
| | Untreated | 24 hr | FC 24hr/0hr | 0 hr | 24 hr | FC 24hr/0 hr |
| Fraction of total mRNA | 3.4% | 3.4% | 1 | 2.6% | 3.3% | 1.27 |
| Fraction of total RPF | 0.4% | 0.7% | 1.75 | 0.5% | 0.6% | 1.2 |

Figure S4

RNA-seq and Ribo-seq datasets of untreated (0 hr) and 24 hr cytokine-treated WT astrocytes (**A**, top) and Mut astrocytes (**B**, bottom) were used to generate relative gene-expression maps (termed 'Proteomaps') (Liebermeister, 2014). A polygon Highlighted in red represents the percentage of transcripts encoding ribosomal proteins (N=77) out of total mRNA (RNA-seq) and total RPF (Ribo-seq) datasets. Polygon size was quantified as described in Methods. (**C**) Fold-Change (FC) differences due to the cytokine treatment in WT and Mut.

### Figure S5

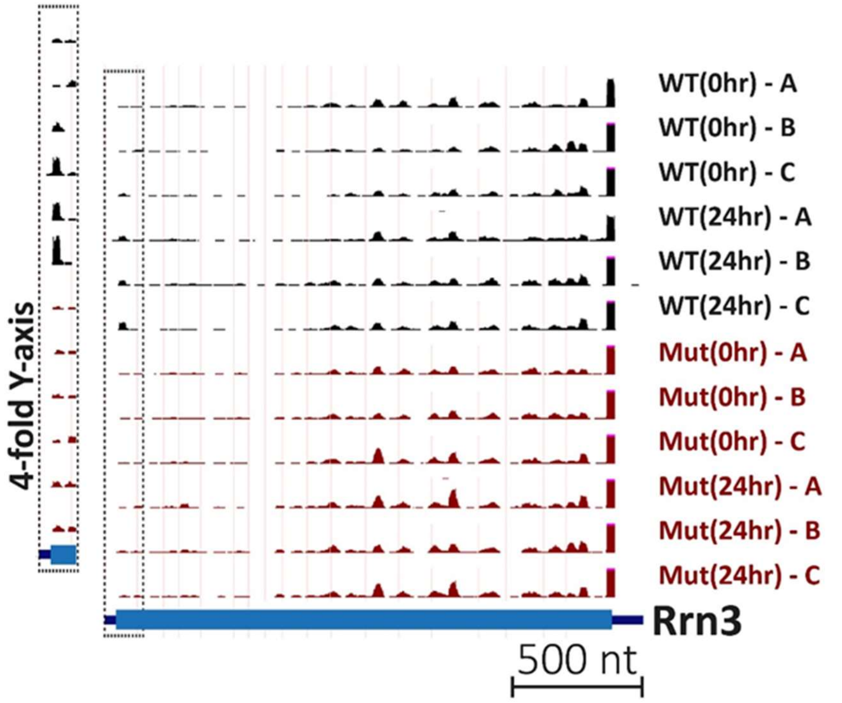


Figure S5

RPF map of Rrn3, highlighting the increase in RPF reads at the AUG start codon, in WT but not in Mut, in response to 24 hr of cytokine treatment (dashed rectangle).

#### Figure S6

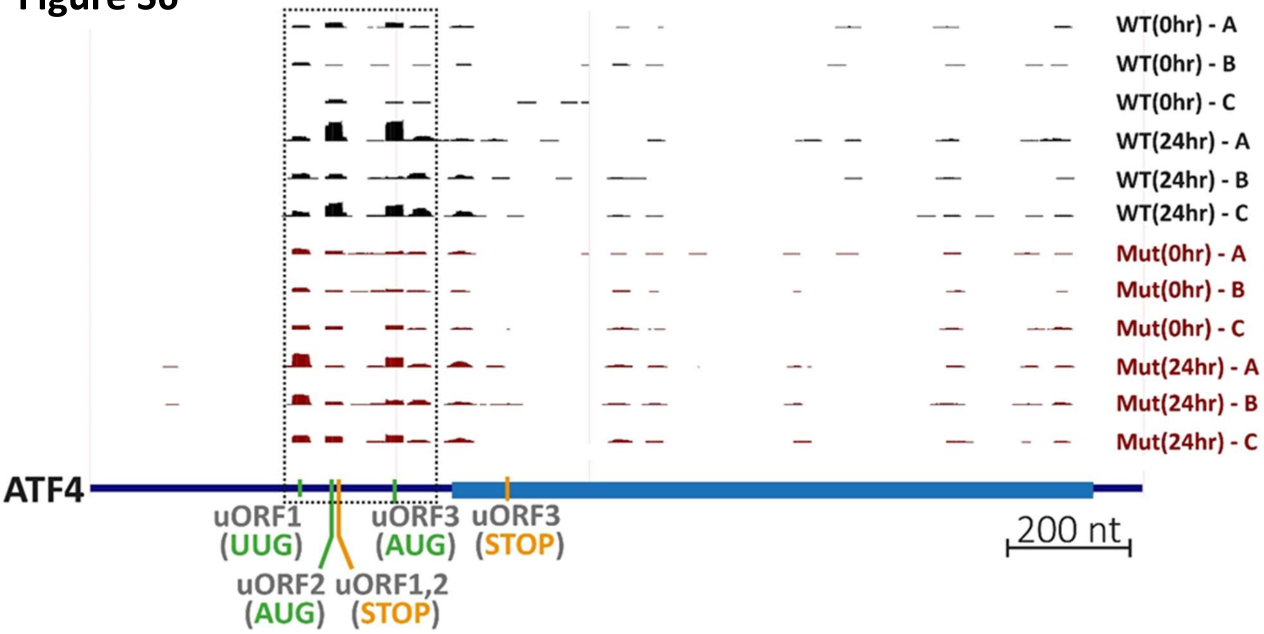


Figure S6

RPF map of ATF4, highlighting the increase in RPF reads at the UUG uORF1 in response to 24 hr of cytokine treatment, more in Mut compared to WT.

5'UTR, dashed rectangle; uORF1 UUG, uORF2,3 AUG, green; uORF1,2,3 stop codons, orange.

#### Figure S7

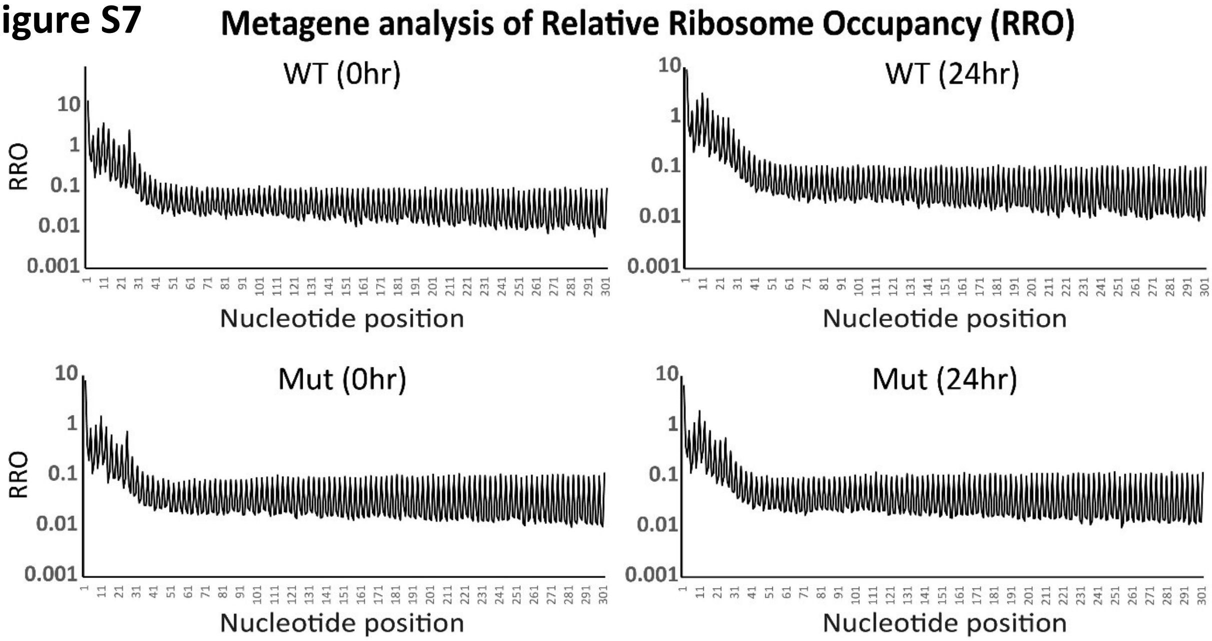


Figure S7

Metagene Analysis of Relative Ribosome Occupancy (RRO). Shown are ribosome protected footprints (RPFs) relative density along the first 300 nucleotides of all the main open reading frames in each of the four Ribo-seq datasets, as indicated. See methods for calculation. A single most abundant transcript was selected per gene. Note the consistent coverage in all datasets and the 'ramp' phenomenon shown at the first 30 nucleotides, as reported by Tuller at al., Cell (2010) 141(2):344-354.
